# Cracking the egg: Isolation and discovery of distinct nonvesicular extracellular particles from *Schistosoma mansoni* eggs

**DOI:** 10.1101/2025.03.24.645107

**Authors:** Madeleine J. Rogers, Athena Andreosso, Jagan Billakanti, Mary G. Duke, Yan Lu, Natasha Collinson, Jian Tan, Laurence Macia, Na’ama Koifman, Matthias Floetenmeyer, Malcolm K. Jones, Catherine A. Gordon, Severine Navarro

## Abstract

Extracellular particles (EPs) derived from helminths are emerging as promising immunomodulatory agents, yet their heterogeneity challenges reproducible purification and characterisation. This study focused on purifying and characterising extracellular particles (EPs) from *Schistosoma mansoni* eggs using tangential flow filtration (TFF) and differential ultracentrifugation (dUC). Contrary to expectations, cryogenic transmission electron microscopy (cryoTEM) revealed that egg-derived EPs are nonvesicular (5–30 nm) nanoparticles (eggNPs) rather than membrane-bound extracellular vesicles. TFF demonstrated superior performance over dUC, yielding higher recovery (76.19% vs. 38.90%), greater purity, and lower endotoxin contamination. Despite limitations in standard protein quantification methods, enriched protein profiles and detection of *S. mansoni* tetraspanin-2 in TFF-purified eggNPs highlights their distinct molecular signatures. The study also identified contaminants in cryopreservation buffers, underscoring the need for optimised storage conditions. Compared to dUC, TFF preserved particle integrity, with cryoTEM images showing fewer aggregates and distinct EggNP morphology. This work proposes TFF as a scalable and reproducible method for isolating *S. mansoni*-derived eggNPs, providing a foundation for their exploration as immunomodulatory agents. These findings not only expand the understanding of helminth EPs but also emphasise their therapeutic potential, particularly in inflammatory and immune-dysregulation disorders.

## Introduction

Extracellular vesicles (EVs) are a heterogenous family of lipid-bilayer membrane-enclosed extracellular particles (EPs) that transport functional and pathogenic cargo, including proteins, nucleic acids, metabolites, and lipids between cells ^1^. Despite our growing knowledge of EVs, their heterogenous nature makes reproducible purification and characterisation complex. In fact, two novel populations of nonvesicular extracellular particles (NVEPs) known as exomeres (<50) and supermeres (<50nm) were recently discovered ^2–4^. These distinct NVEPs, like EVs, contain immunomodulatory cargo that play essential roles in immune modulation, angiogenesis, and disease progression ^5–9^. The diversity of EPs is such that they provide exciting avenues for diagnostic/prognostic biomarkers and have the potential for therapeutic translation into novel treatments for cancer, parasitic diseases, and immune dysregulation disorders.

Like other organisms, helminth EVs exhibit exquisite cell and tissue specific functions ^10–12^. This is necessitated by their complex life cycle within the mammalian host that requires migration through multiple tissues, a process they achieve through the release of various mucosal- and cell-specific compounds, a large number of which are enclosed in EVs ^11,13^. The immunomodulatory potential of schistosome-derived EVs has been well described since their discovery in 2012 ^14^, and have been shown to promote immune tolerance in a myriad of inflammatory disease models including inflammatory bowel disease and allergic disease (extensively reviewed elsewhere) ^15^. This is due to their ability to promote regulatory T cells and B cells, making schistosome EVs of special interest for therapeutic translation ^13,16–19^. There has been extensive research into the isolation and characterisation of EVs from adult schistosomes, with most relying on a combination of differential ultracentrifugation (dUC) using density gradients or precipitation reagents for isolation ^5,16,17,19–30^. The ability is of *Schistosoma* spp. eggs and their excretory/secretory products to promote immunomodulation is well established ^31^. However, research into *Schistosoma* egg-derived EVs is limited, with only four papers published ^32–35^. Furthermore, the methods used in these studies to prove the presence of EVs in isolated samples were not compliant with requirements outlined in the Minimal information for studies of extracellular vesicles 2023 (MISEV2023) ^32–34,36,37^. To date, there are no studies focussing on schistosome-derived NVEPs; illustrating a previously overlooked novel source for therapeutic translation.

To effectively characterise molecular composition and function of EPs, it is essential to purify them of large molecular contaminants that may exhibit similar size or buoyancy. Helminth EPs can also become contaminated by host, bacterial, and parasite residues ^38^. With each purification step, EPs are lost due to their rupture or absorption onto plastic surfaces during differential ultracentrifugation (dUC), limitations that are yet to be characterised for NVEPs ^39–41^. While the use of larger volumes of starting material can compensate for loss, the challenge with helminths is the limited access to the parasite material required for the production of sufficient EPs ^42^. This is a particular issue with *Schistosoma* spp. due to their complex life cycle that requires access to definitive mammalian and intermediate snail hosts. In addition, *S. mansoni* has relatively low egg output, resulting in lower yields than that of *S. japonicum* infections ^43^. Obtaining relevant schistosome life cycle stages for culture is limited by mammal-infecting larvae (termed cercariae) and the number of parasites that can be collected from mammalian hosts. Schistosomula (cercariae that have transformed after invasion of host skin) can be *in vitro* culture or from the skin and lungs of infected mammalian hosts. Worms are collected by portal vein perfusion, and eggs are isolated from enzymatically digested infected liver and gut tissues. Miracidia, the intra-ovular larvae, can easily be hatched from eggs, however, to date there has been no attempt to isolate EPs from this stage, likely due to their lack of relevance to mammalian hosts. In addition, the requirement of fresh water for miracidial hatching complicates the study of EPs, which require specialised buffer compositions to preserve structural and/or membrane integrity ^44,45^.

Other EV purification methods include antibody-based bead-capture technologies, precipitation reagents such as ExoQuick^TM^, polyethylene glycol (PEG), and immunoaffinity capture-based techniques. These methods often result in the co-precipitation of contaminating proteins, RNA, and circulating DNA, and can alter the biological activity of the EVs ^46–48^. Therefore, the above methods are often used in combination with UC to improve yield and purity ^49^. Size exclusion chromatography (SEC) has shown to be a promising technique for producing high yield helminth EVs with a higher purity ^50^. However, the requirement of such large starting sample volume and the complexity of contaminant profiles associated with schistosome egg-derived EPs means that these protocols continue to pose significant challenges to effective purification in terms of purity, yield, and overall processing costs. More recently, novel approaches such as tangential flow filtration (TFF) show promising results for faster EV separation from larger volumes ^51^. Combining TFF with multimodal chromatography resin-based approaches can further purified the TFF concentrated EPs ^52,53^. Asymmetric-flow field-flow fractionation (AF4) technology has also been used successfully to separate exomeres from EVs for subsequent molecular and structural characterisation ^2,3^. Despite these recent developments, there is significant divergence observed in helminth EP structure and function compared to their mammalian counterparts ^12^. Additionally, most available antibody markers are not specific to schistosome egg-derived EPs. Together these pose unique challenges for reproducible isolation and purification of high purity EPs from helminths for *in vitro* and *in vivo* studies.

The potential for schistosome EPs and their protein/miRNA cargo for immunomodulation makes them exciting prospects for therapeutic development. Good Laboratory Practice (GLP) and the appropriate biosafety control measures are essential to ensure samples are highly pure and do not contain contaminating products (proteins, nucleic acids, metabolites, bacterial endotoxins, etc.) for studies. This is particularly important for therapeutic translation, for example, for the discovery of vaccine candidates for schistosomiasis ^20,33^. Therefore, this study aimed to develop a scalable method for the isolation of EVs from *S. mansoni* eggs using tangential flow filtration (TFF) method, which allows the concentration and separation of highly pure particles from large (up to 20 L) starting volumes while reducing handling time. To determine the viability of TFF, EPs were also purified using differential centrifugation (dUC) as a comparator.

## Materials and methods

### In vitro cultivation of S. mansoni eggs

#### Sample collection and processing

Seventy-five (7-9-week-old) Swiss outbred female mice were infected percutaneously with 150-190 cercariae from *S. mansoni*. *Biomphalaria glabrata* snails infected with *S. mansoni* were provided by the NIAID Schistosomiasis Resource Center of the Biomedical Research Institute (Rockville, MD) through NIH-NIAID Contract HHSN272201700014I for distribution through BEI Resources. When the infection reached patency (approx. 5-6 weeks post-infection), animals were injected with Heparin, an anticoagulant (100 µL intraperitoneal/mouse). After CO_2_ (2.2L/minute) euthanasia, the hepatic portal vein was perfused to remove blood and adult worms, and egg-infected livers were collected from the mice. Live *S. mansoni* eggs were purified from the livers of infected Swiss mice using a collagenase B digestion as previously described ^54^. In addition, a number of infected livers were collected and stored at −20□ for soluble egg antigen purification.

#### Culture of eggs and tracking of survival

The isolated eggs were counted under a light microscope and egg viability was determined (**Day 0**) before culturing by staining with 0.4% Trypan blue, a vital stain that stains dead or cracked eggs blue. Isolated eggs were then cultured in flasks at densities of 1000-1500 egg/mL in Roswell Park Memorial Institute (RMPI) 1640 GlutaMAX medium (ThermoFisher, 61870127) containing 100 U/mL Penicillin and 100 µg/mL Streptomycin (P/S) (ThermoFisher, 15140122). The eggs were kept at 37°C in a humidified incubator with 5% CO_2_. Every 22-24 hours (**Day 1, 2, and 3**), the contents were taken, centrifuged gently at 800 *g* for 5 mins to separate eggs from egg conditioned media (ECM), which were placed back into culture flasks with fresh media. Egg viability was assessed at each time point by staining 500-1000 eggs with Trypan blue as per day 0.

ECM was then filtered (0.2 µm aPES membrane) and centrifuged at 3500 *g* for 30 mins at 4°C to remove pellet and remove cellular debris. The resultant clarified ECM (cECM), which contained all egg-derived EPs, was separated into 50 mL Falcon tubes containing 20-30 mL cECM, snap frozen in liquid N_2_ to preserve membrane integrity, and stored at −80°C until sufficient material was collected for EP isolation and purification.

#### Preparation of S. mansoni soluble egg antigen (SEA)

*S. mansoni*-infected livers were thawed from −20°C storage on ice and eggs were purified using collagenase B digestion as previously described ^54^. Purified eggs were resuspended in 750 µL 1 mM PBS, counted using compound microscopy, and separated evenly into groups of approximately 250,000 eggs/mL in three microcentrifuge that contained approximately 68,500 eggs. The eggs in each tube were then ruptured using a Kimble® handheld homogeniser (DWK Life Sciences, SCERSP749540-0000) and disposable micro pestles (Roch, CXH7.1) for three to four bursts of 60 seconds. A small aliquot (10 µL) of resuspended material was taken from each tube and examined by compound microscopy to confirm that approximately 95% of eggs were disrupted. Samples were then centrifuged at 200 x *g* for 20 minutes at 4°C and the supernatant was collected and ultracentrifuged for 90 minutes at 100,000 x *g* and 4°C using a Hitachi himac S55A2 rotor in a Hitachi himac CS150X benchtop micro ultracentrifuge. The supernatant was then collected and sterilised by passing it through a 0.2 µm syringe filter. All samples were stored at −20°C until analysis.

### Isolation and separation of *S. mansoni* egg-derived extracellular particles

#### Tangential Flow Filtration (TFF)

Two biological replicates were completed for eggNP isolation. cECM or control media was thawed overnight at 4°C before isolation and pooling. The same protocol was also undertaken on control media (RMPI containing 1% P/S), 1 mM PBS, and 1 x a cryopreservation buffer (1 mM PBS, 1 mM PBS, 5% sucrose, 50 mM Tris, 2 mM MgCl_2_). To enrich, isolate, and purify EPs, concentration and buffer exchange was undertaken using a MidGee Hoop tangential flow ultrafiltration hollow fibre cartridge with a 300 kDa cut-off (Cytiva, UFP-300-E-H42LA). Using an AKTA Flux S tangential flow filtration system (Cytiva), conditioned media was concentrated through the hollow fibre cartridge following material equilibration with the 1 mM PBS at a maximum shear rate of 2000 s^-1^ to protect EP integrity. This generated two fractions, the retentate containing EPs and some contaminants that are larger than 300 kDa, and the permeate containing non-EP contaminants that are smaller than 300 kDa. The permeate was snap frozen in liquid N_2_ and store for further analysis at −80□. The EP-containing retentate was then washed with 1 mM PBS containing 1.5 M NaCl (pH 7.4) by stepwise addition and recirculation through the hollow fibre cartridge to remove additional impurities by breaking electrostatic interactions, which was continued for six retentate volumes (30 mL) of buffer exchange. Subsequently, the salt was removed by stepwise addition and recirculation of six retentate volumes of (30 mL) 1mM PBS. Two aliquots of 30 µL were taken for CryoTEM imaging and characterisation, then a final buffer exchange step was undertaken using six retentate volumes (25 mL) of EV cryopreservation buffer (1 mM PBS, 5% sucrose, 50 mM Tris, 2 mM MgCl_2_). The final retentate (15 mL), which was 160 times concentrated, was split into 14 aliquots of 1 mL and 2 aliquots of 500 µL before being snap frozen in liquid N_2_ and stored at −80°C until further analysis.

#### Differential ultracentrifugation (dUC)

*S. mansoni* egg-derived EPs were isolated using dUC as previously described with some alterations ^32^. Firstly, cECM was thawed overnight at 4□ then placed into polycarbonate ultracentrifuge (26.3 mL) bottles, closed with the aluminium cap assembly (Beckman Coulter 355618), and ultracentrifuged in batches at 10,000 *g* for 60 min at 4□ in a Beckman Coulter Optima L100XP floor standing ultracentrifuge using a Type 50.2 Ti rotor (Beckman Coulter, 337901). The supernatant was collected and ultracentrifuged at 120,000 *g* for 4 hours at 4□ in the same rotor and ultracentrifuge. The pelleted EPs were then resuspended in 10 mL 1 mM PBS, placed into 1.5 mL polypropylene tubes, and ultracentrifuged under the same conditions for 4 hours in a Hitachi himac CS150X benchtop micro ultracentrifuge using a Hitachi himac S55A2 rotor. The EP pellet was resuspended in 5 mL EV cryopreservation buffer (1 mM PBS, 5% sucrose, 50 mM Tris, 2 mM MgCl_2_, pH 7.4), and the final concentrated EP sample was split into aliquots of 500 µl, snap-frozen in liquid N_2,_ and stored at −80□ until analysis.

### Characterisation of *S. mansoni* egg-derived EPs

#### Negative staining transmission electron microscopy (NS-TEM)

The eggNPs prepared from each isolation protocol were imaged by negative staining transmission electron microscopy (TEM) as previously described [242]. Briefly, 10 µL of purified eggNP suspension was applied to a 300-mesh copper carbon-formvar coated grid (ProSciTech, GSSFC300CU-SB) and incubated for 2 mins to allow attachment. The grids were then blotted dry using filter paper, negatively stained using 10 µL of 1% uranyl acetate for 2 min and allowed to dry. The eggNPs were observed on a Hitachi HT7700 transmission electron microscope available at the University of Queensland (UQ) Centre for Microscopy and Microanalysis (CMM) operating at 80 kV.

#### Cryogenic transmission electron microscopy (CryoTEM)

Carbon-formvar coated lacey EM grids (200 mesh; LFC 200-Cu-50) were glow-discharged by 0.11 mbar air for 4 minutes using the low pressure plasma system (Diener Electronic Pico PCCE). EggNP preparations (in 1mM PBS) from *S. mansoni* eggs were prepared in an EM GP2 robotic vitrification system (Leica Microsystems, Germany) at a controlled temperature (22°C) and humidity (95%). Three microlitre of eggNPs was placed on a glow-discharged grid and excess suspension was automatically removed by blotting for three seconds on Whatman #1 filter paper. The grid was directly plunged into liquid ethane at its freezing point (−183°C). Vitrified grids were then rapidly transferred into liquid nitrogen for storage. Samples were imaged at −176°C in a frozen hydrated state using a CryoARM 200 (JEOL, JEM-Z200FSC) TEM located at the University of Queensland Centre of Microscopy and Microanalysis. This microscope is equipped with a cold-field emission gun (FEG), and an in-column Omega energy filter. Zero energy loss images were acquired at an accelerating voltage of 200 kV and a filter setting of 20eV. A Gatan K2 direct detector camera was used under low-dose conditions using SerialEM software ^55^ to record all images.

#### Measurement of particle size using Fiji

The particle diameter of *S. mansoni* egg-derived EPs were measured from cryoTEM images using Fiji Is Just ImageJ (Fiji) (National Institutes of Health, USA.; ImageJ v. 1.). Attempts were made to analyse eggNP size using the Nanodefine particle sizer plugin v. 1.0.7. However, sufficient contrast could not be achieved for the Nanodefine plugin to be effective due to the significant size and shape variation, aggregation of eggNPs, and the presence of contaminating micelles in cryobuffer. Therefore, particle diameter (nm) was manually measured in Fiji after the scale was set using the “line tool” followed by selecting “Measure” in the Analyze menu or pressing “ctrl M” on the keyboard. For each purification protocol, 104 particles were measured across five cryoTEM images.

#### Nanoparticle tracking analysis

The EP particle size and concentration were assessed by NTA on a Nanosight NS300 (Malvern Instruments Limited) and the ZetaView (Particle Metrix). For measurement on the Nanosight NS300, samples were diluted in 0.22 µm filtered PBS so that an optimal 20-100 particles per frame was captured. Three 60 second videos were acquired with the syringe pump speed set at 50 µl/s and temperature at 25°C. The NTA 3.2 software version was used to calculate total concentration and size distribution. For measurement on the ZetaView, samples were diluted in 0.22 µm filtered PBS so that an optimal number of particles (50-200 particles in view) were detected at the optimal sensitivity setting (80-85). For analysis, 3 acquisitions per samples of 1 cycle and 11 positions were recorded to calculate total particle concentration and size distribution. For both machines, the same settings were used for running all samples within the same experiments.

### Protein quantification and characterisation

#### Protein quantification assays

Protein quantification was undertaken using the Qubit^TM^ Protein Assay Kit (Thermo Fisher Scientific, Q33212) and MicroBCA^TM^ (Thermo Fisher Scientific, 23235) according to the manufacturers protocol. The Qubit assay kit includes 3 standards for calibration. 5 - 20 µL sample was diluted in 180 – 195 µL Qubit^TM^ protein reagent A and protein buffer B (1:199) in thin wall PCR tubes (Thermo Fisher Scientific). The concentration of each was read using a Qubit^TM^ 4 Fluorometer (Thermo Fisher Scientific). For microBCA, a standard curve was prepared using Bovine Serum Albumin, with final concentrations in µg/mL of 2000, 1000, 500, 250, 125, 63, 31, 16, 8, 4 and 0. For BCA, a standard curve was prepared using Bovine Serum Albumin, with final concentrations in mg/mL of 2, 1, 0.5, 0.25, 0.125, and 0. EggNP samples were diluted 1:8, 1:16, or 1:30 while SEA was diluted 1:2, 1:4, 1:8, and 1:16 in 1 mM PBS before analysis. Protein quantification was also undertaken with a NanoDrop^TM^ 2000 UV-Vis spectrophotometer (Thermo Fisher Scientific) using 1 µL sample, which were measured in triplicate.

#### Determination of particle purity

The purity of EPs was determined in accordance with recommendations by Webber and Clayton (2013) using the ratio of particles/µg of protein with the following equation ^56^. Particle purity was calculated using all Qubit^TM^ protein quantification assays.

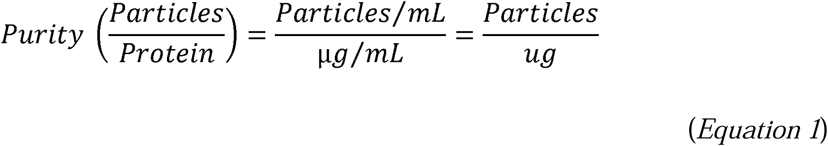

We also calculated the ratio of protein (ng) per *S. mansoni* egg as an additional protein-based purity measure with the following equation.

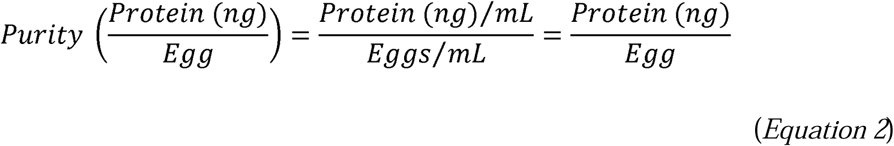

Finally, due to variability in protein quantities, the purity of EPs was also measured using the ratio of particles per *S. mansoni* egg with the following equation.

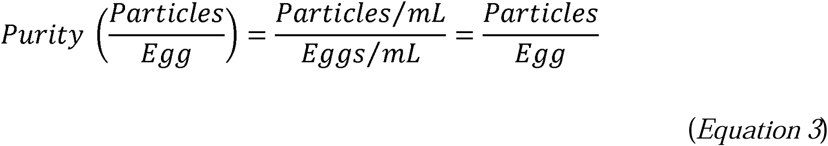

#### SDS-PAGE

Samples were resolved by SDS-PAGE to visualise their protein profile on 4-20% Mini-PROTEAN TGX precast polyacrylamide protein gels (Bio-Rad, 4561094) using the Precision Plus Dual Xtra Prestained Ladder (Bio-Rad, 1610377). *S. mansoni* SEA was purified and 1 – 10 µg was combined with 4 x MES (2-(N-mopholino)-ethanesulfonic acid) buffer then reduced and denatured at 75°C for 5 min. Gels were stained using Coomassie Brilliant blue R-250 (Merck, 112553) for one hour and destained overnight in destaining solution (40% v/v methanol, 10% v/v acetic acid, 50% v/v H_2_O). For silver stain, 0.5 - 1 µg of each sample was denatured with 4 x MES as above and gels at were stained using Bio-Rad Silver Stain Plus^TM^ Kit (Bio-Rad, 1610449) following the manufacturer’s instructions for 3-20 minutes. Gels were then visualised using the Gel Doc EZ Imager (Bio-Rad, Australia).

#### Western blotting

Samples were thawed on ice and resolved by SDS-PAGE as outlined above. The protein bands were subsequently transferred onto methanol activated nitrocellulose membranes, which were blocked with 5% skim milk (w/v) in 1 mM PBS supplemented with 0.1% (v/v) Tween (1 mM PBST) for one hour. The blocked membranes were then incubated overnight at 4°C with the following primary antibodies: anti-HSP90 (Cell Signalling, 4877), which was used in a 1:1000 dilution, and rabbit polyclonal sera against tetraspanin-2/5B, a conserved region of a hookworm aspartic protease that recognises *S. mansoni* (*Sm*)TSP-2, which was used in a 1:2500 dilution (kindly provided by Professor Alex Loukas and Dr Darren Pickering, James Cook University, Australia). Incubated membranes were washed three times for 15 min in 1 mM PBST, incubated for one hour with horseradish peroxidase (HRP)-linked Goat anti-Rabbit IgG (Sigma 12-348) or anti-mouse (Sigma 12-349) secondary antibody (1:3000) and washed in 1mM PBST again as above. SuperSignal^TM^ West Pico PLUS chemiluminescence substrate (ThermoFisher, 34580) was applied to each membrane following the manufacturer’s instructions and an iBRIGHT CL1500 Imaging System (ThermoFisher, Brisbane, Australia).

#### Statistical analysis

Statistical analysis was completed using Graph Prism v. 9.5.1 (GraphPad Software, San Diego, CA, USA). For results obtained by, Qubit, microBCA, nanodrop, and NTA, groups were compared using a one-way analysis of variance (ANOVA) followed by a Tukey’s multiple comparisons test. Particle diameter measurements were performed using Fiji and were compared using a Welch’s two-tailed t test or a one-way ANOVA and Tukey’s test. Data from all experiments are expressed as means ± standard deviation (SD).

## Results

### *S. mansoni* egg survival during *in vitro* culture

*S. mansoni* eggs were isolated and purified from the livers of infected mice as shown in **Figure 1**. The isolated eggs were analysed under a light microscope and counted, and we observed that the majority of eggs were mature, with a visible miracidium that exhibited movement (**Figure 2A)**. Egg viability was also determined (**Day 0**) before culturing by staining with 0.4% Trypan blue, a vital stain that stains dead or cracked eggs blue (**Figure 2B)** ^57^. Isolated eggs were then cultured in cell culture flasks at densities of 1000 egg/mL in serum-free RMPI-1640 media containing 1% P/S. Every 22-24 hours when conditioned media was collected and replenished, 500-1000 eggs were collected, and morphology and viability were assessed (Day 1 – 3) (**Figure 2C**).

**Figure 1:**
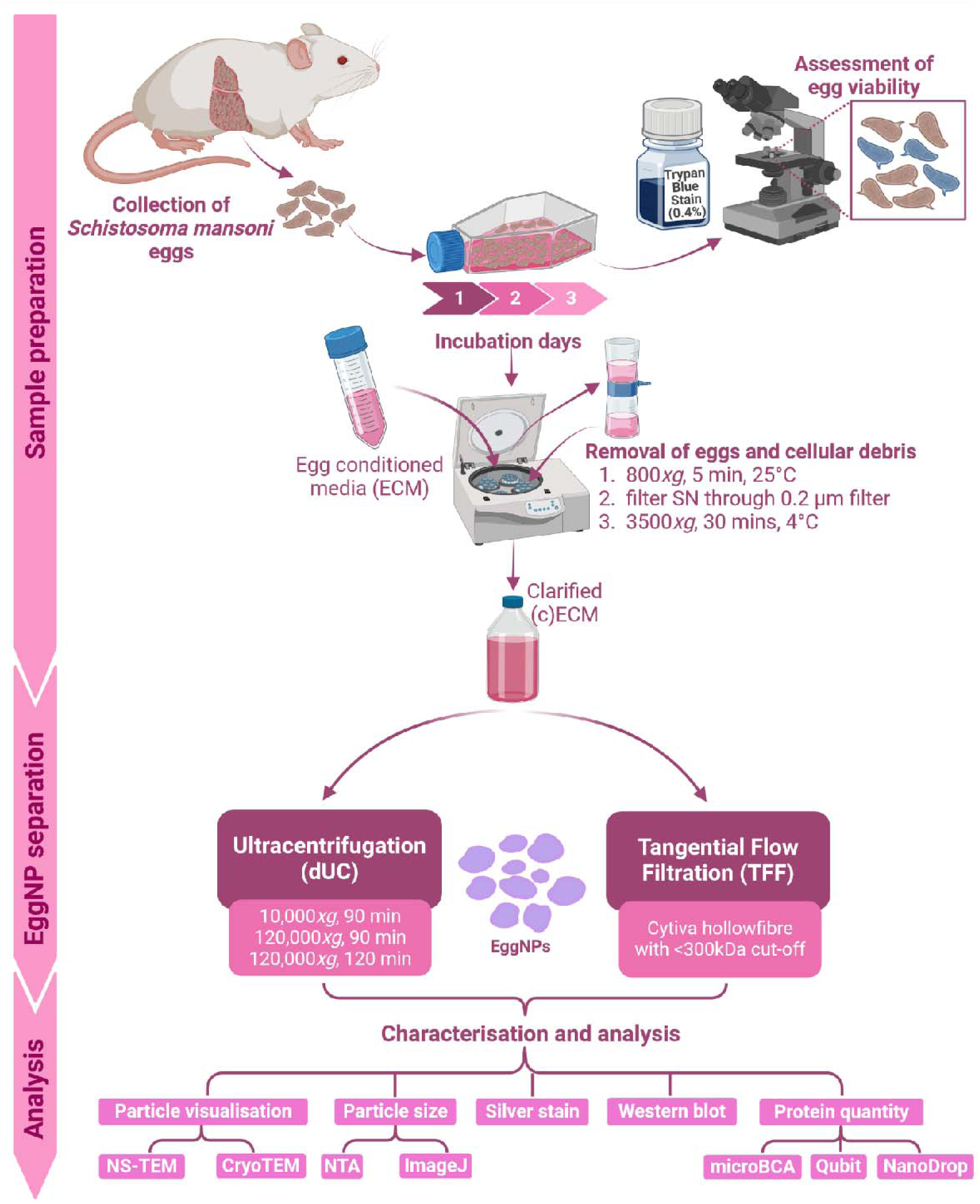
Schematic summary of study design. *Schistosoma mansoni* eggs were isolated from infected mouse livers and cultured for three days and egg survival was tracked every 24 hours with Trypan blue. Egg conditioned media (ECM) was collected and cleared of eggs and cellular debris by centrifugation to give clarified (c)ECM. EggNPs were isolated and purified from cECM by differential ultracentrifugation (dUC) or tangential flow filtration (TFF). EggNPs were characterised visually by negative staining transmission electron microscopy (NS-TEM) and cryogenic TEM, sized using nanoparticle tracking analysis (NTA) and ImageJ, and cargo was characterised using silver stain (SDS-PAGE) and Western blot. Finally, protein content was quantified by Nanodrop, microBCA, and Qubit. Created with BioRender.com

**Figure 2:**
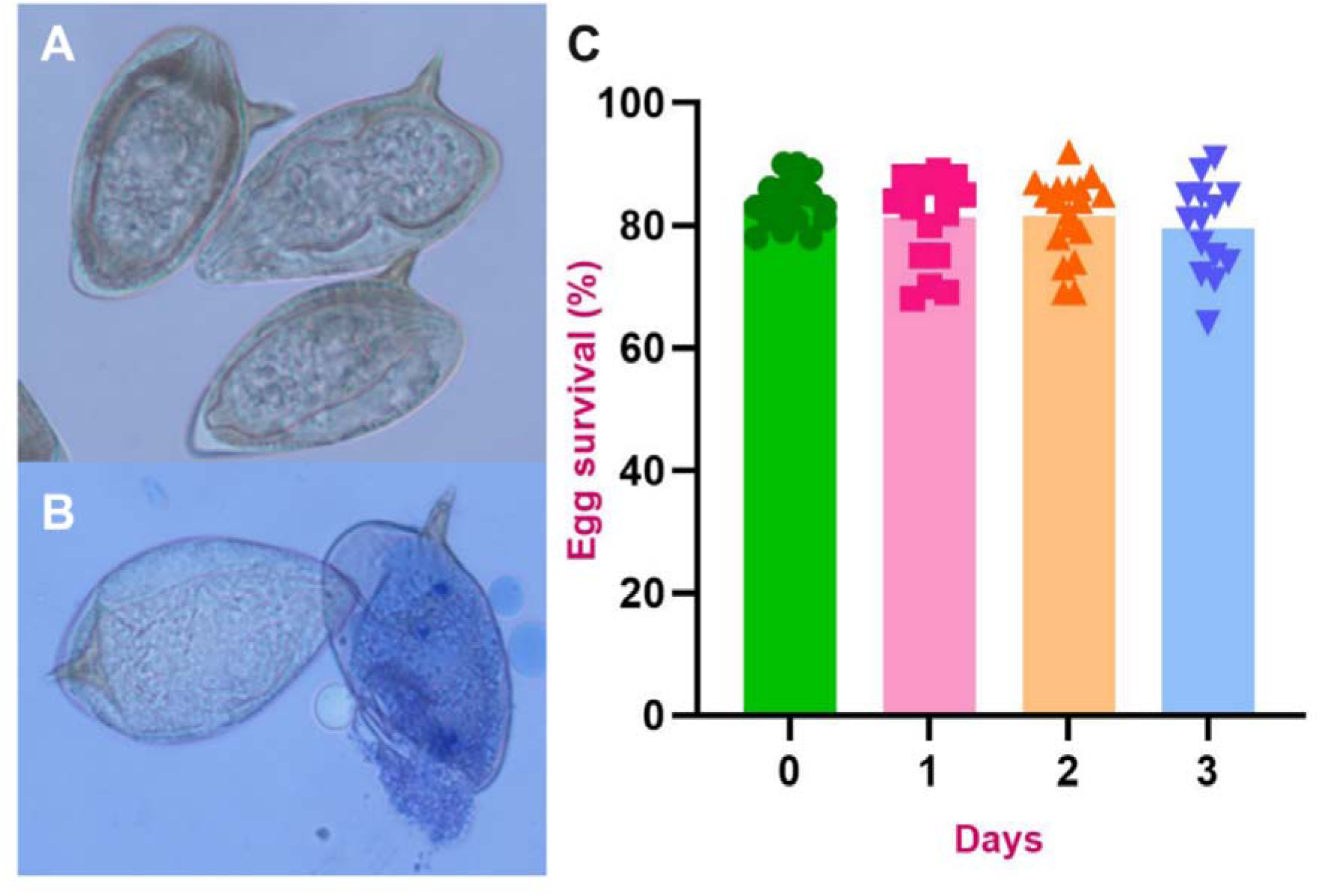
*Schistosoma mansoni* egg viability during culture. (A) Mature *S. mansoni* egg with developing miracidia enclosed in a thick protein shell. (B) Viable *S. mansoni* egg next to unviable cracked egg and miracidia stained with 0.4% Trypan Blue vital stain. (C) *S. mansoni* egg viability following isolation (Day 0) and during 3 days of culture in RMPI-1640 supplemented with 1% PenStrep. Results are shown as mean ± SD (one-way ANOVA with Tukey’s multiple comparison test).

Egg morphology did not vary significantly across days and miracidia remained active in viable eggs, with the majority of unviable eggs being present due to the eggs hatching. Although schistosome eggs will usually only hatch in freshwater, we observed that a small number of eggs hatched during culture. Egg survival remained relatively constant across three days of culture, with no significant difference observed in viability between each day. However, the variability of counts increased with each day (**Figure 2C)**. In addition, it was observed that the eggs became progressively more aggregated with each day and became stuck to the sides and bottom of the culture flask. *S. mansoni* eggs contain large amounts of adhesive compounds, such as carbohydrates, which are either secreted or released when the eggs hatch ^58^. In addition, we observed (using compound microscopy) that a large number of eggs, particularly the shells from hatched eggs, remained stuck to the walls of the flasks. Therefore, it is possible that egg viability decreased further over each day, but most unviable eggs became stuck to the flask and could not be counted. The minimal changes in egg viability over the three days allowed for cECM from all three days to be pooled for further analysis.

### *S. mansoni* eggs produce only nonvesicular extracellular nanoparticles (EggNPs) in culture

*S. mansoni* egg-derived EPs were isolated and separated from serum-free cECM using either dUC or TFF. NTA and NS-TEM indicated a significantly more diverse nanoparticle size range (20-150 nm) than originally hypothesised (Data not shown). This necessitated a change to using hollow-fibres with a lower molecular weight cut-off of 300 kDa instead of the initial 500 kDa cut-off to prevent loss of the smaller particles.

Visualisation of dUC, TFF + dUC isolated eggNPs by NS-TEM and NTA suggested the presence of nanovesicles with similar size ranges to EVs. However, extensive imaging by cryoTEM of all preparations showed no evidence of membrane-bound vesicles (**Figure 3**). However, a population of previously uncharacterised nonvesicular extracellular particles (NVEPs) was discovered. The same NVEPs were not observed in TFF preparations from egg-free media or buffers, indicating they are egg-derived (EggNPs). Due to the consistent issues experienced with imaging EggNPs by NS-TEM (**Figure S1**), all subsequent preparations were imaging by cryoTEM only. EggNPs have an irregular shape, are 5-30nm in diameter, and aggregate significantly (**Figure 3)**. EggNPs separated by dUC appeared to aggregate more closely (**Figure 3A-B**) compared TFF eggNPs (**Figure 3D-E**). Particle diameter of eggNPs was measured using Fiji and eggNPs isolated by dUC (**Figure 3C**) were found to have an average of 17.09 ± 4.60 nm, while TFF separated eggNPs (**Figure 3F**) were found to have a significantly smaller diameter of 13.49 ± 2.52 nm (p<0.0001, paired two tailed t test). Small amounts of contaminants were present in all TFF separated eggNP samples stored in 1mM PBS in the form of aggregated microtubule-like structures (10 nm wide and 30-100 nm long), which were not observed in eggNPs stored in cryobuffer (**Figure 4A-B**). In cryobuffer-stored eggNPs, we observed organised filaments (100 nm wide and over 5 µm long) embedded with eggNPs that exhibited periodicity (**Figure 4C-D).** Smaller filaments of similar length (over 5 µm) were also observed, which are likely disorganised forms of the above cross-hatching filaments (**Figure 4E-F**). These structures likely originate from cilia (the microtubule-like structures) found on miracidia and the outer envelope of eggs that hatch during culture (the fibrils), which also explains their relatively low abundance as only small numbers of eggs hatched during culture. These were not observed in dUC isolated eggNPs samples, most likely because they would pellet when the samples were centrifuged at 10,000 *g* to remove large contaminants.

**Figure 3:**
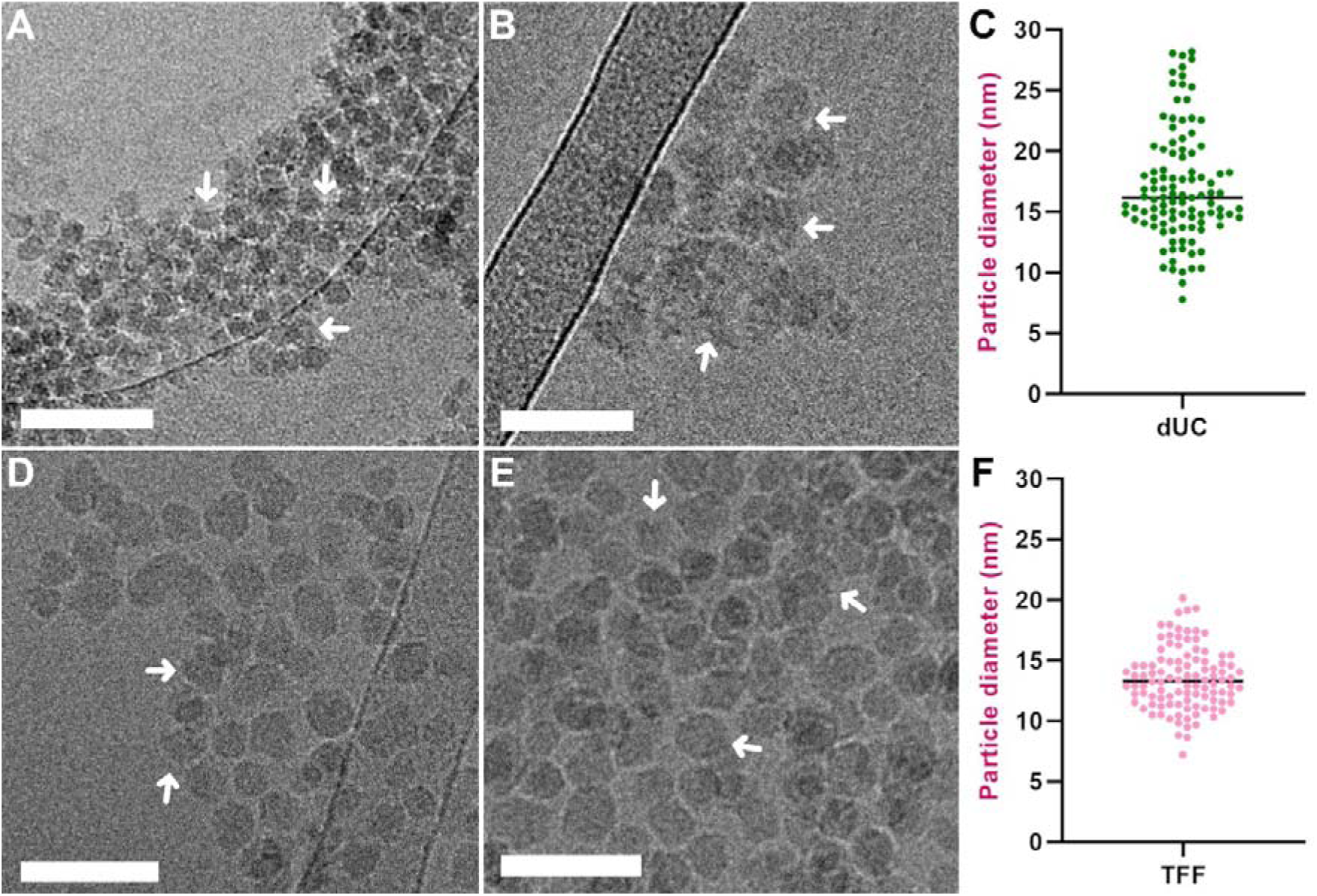
Cryogenic transmission electron microscope micrographs and diameter analysis of *Schistosoma mansoni* nonvesicular extracellular particles (eggNPs). Differential ultracentrifugation (dUC) separated eggNPs imaged by cryogenic transmission electron microscopy (TEM) at 60,000x magnification showing significant aggregation (A) and distinct rosette of tightly aggregated eggNPs (B). (C) ImageJ analysis of dUC separated eggNP diameter = 17.09 ± 4.60 nm. (D-E) Tangential flow filtration (TFF) separated eggNPs imaged by cryoTEM at 60,000x magnification. (E) ImageJ analysis of TFF eggNP diameter = 13.49 ± 2.52 nm. Scales = 50 nm. Diameter of each separation method was compared by Welch’s two-tailed t test, finding a significant difference (p<0.0001). Data are represented as mean ± SD. n = 104 particles were measured over five cryoTEM images for each separation protocol.

**Figure 4:**
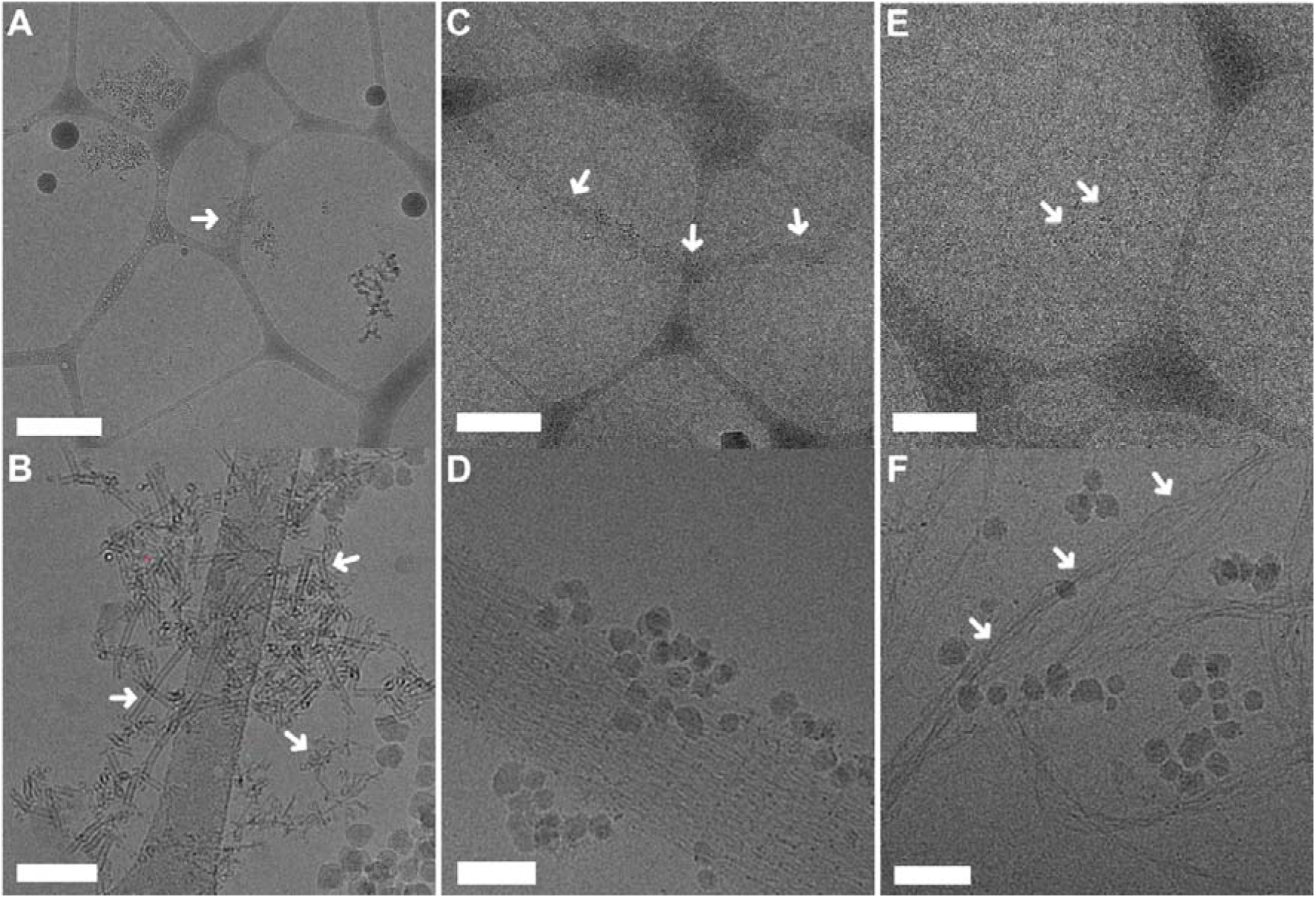
Cryogenic transmission electron microscope micrographs of contaminants found in tangential flow filtrations (TFF) separated *Schistosoma mansoni* nonvesicular extracellular particles (eggNPs). (A) *S. mansoni* TFF eggNPs and microtubule-like contaminants in 1mM PBS at 6000x or (B) 60,000x magnification. (C) Organised fibrillar contaminants with embedded eggNPs at 6,000x (D) or 60,000x magnification. (E) TFF eggNPs and unorganised fibril contaminants at 6,000x (F) or 60,000x magnification. Samples in images C-F were stored in cryopreservation buffer. Scale for A, C, and E = 500 nm; Scale for B, D, and F = 50 nm.

### EV cryopreservation buffer contains contaminating sucrose-derived particles

NTA was undertaken on TFF preparations isolated from on 0.22 µm filtered egg free control media which were stored in either 1mM PBS or cryobuffer as discussed above to evaluate for particle contamination. The particle concentration was found to be 1.7 x 10^7^ and 6.2 x 10^9^ particles/mL for media control particles in PBS and cryobuffer respectively. To examine why cryobuffer samples had significantly higher than expected particle concentration we performed cryoTEM on control NP samples in both 1mM PBS and cryobuffer (**Figure 5**). Although the 5% sucrose used in the cryopreservation buffer is meant to preserve membrane and cargo integrity, it seems that this sucrose concentration also results in the formation of contaminating sucrose particles. These were clearly visible in cryoTEM images surrounding the eggNPs and media control preserved in cryobuffer (**Figure 5D-F**), which were not visible around particles stored in PBS (**Figure 5A-C**). These formed a granular layer over the grid, decreasing image contrast (grainy), or could on occasions aggregate around regions of eggNPs, obscuring them. Despite the presence of contaminating sucrose particles in cryobuffer preserved eggNPs, their concentration was negligible compared to that of the eggNPs (**Figure S2**), confirming that all NVEP present in eggNP preparations originate from *S. mansoni* egg. Given that preserving vesicular membranes was no longer necessary, cryobuffer was excluded for subsequent eggNP purifications, which were stored in 1mM PBS only, significantly improving the visibility of eggNPs (**Figure 5A-C)**.

**Figure 5:**
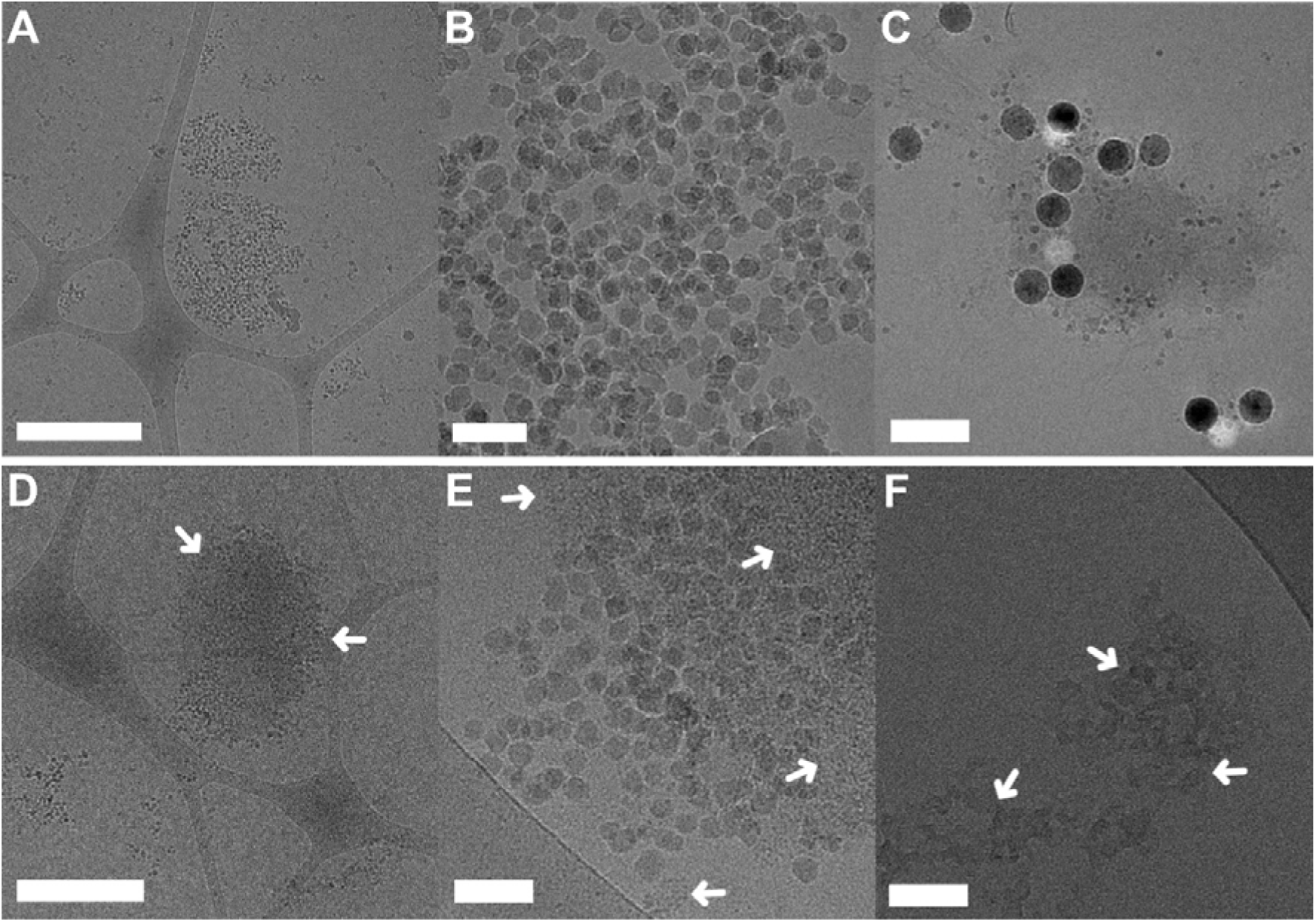
Cryogenic transmission electron microscope micrographs of *Schistosoma mansoni* nonvesicular extracellular particles (eggNPs) or media control samples that were separated using tangential flow filtration (TFF) and stored in either in (A-C) 1mM PBS or (D-F) EV cryopreservation buffer (1mM PBS, 5% (w/v) sucrose, 50mM tris, 2mM MgCl2, pH 7.4). (A) TFF separated eggNPs in 1 mM PBS at 6,000 and (B) 60,000 times magnification. (C) Media control TFF sample (60,000 x). (D) TFF separated eggNPs in cryobuffer at 6000 and (E) 60,000 times magnification. (F) Media control TFF sample (60,000 x magnification). The scales for A and D = 500 nm. Scale for B-C and E-F = 50nm.

### TFF and dUC gave similar particle concentration and different size distribution

EggNP size and concentration from each separation method was analysed by NTA. The size for the starting material (cECM) was found to be 49.6 – 142.3 nm (**Figure 6A**) while dUC eggNPs ranged from 53.4 – 163.8 nm, peaking at 89.5 nm (**Figure 6B**). The TFF enriched eggNPs exhibit a similar size range (51.7 – 162.8 nm), however particle concentration peaked at 95.07 nm (**Figure 6C**). To confirm that these results were not due to the presence of contaminating particles from the media or 1mM x PBS, NTA was undertaken on 1mM PBS and egg-free media (EFM) individually, or TFF enriched EFM with the final buffer exchange into 1mM PBS. The particle distribution from 1mM and EFM was subtracted from the dUC data, while the TFF enriched EFM distribution was subtracted from the TFF data. Results indicated that any particle contamination was non-significant (**Figure S1**).

**Figure 6:**
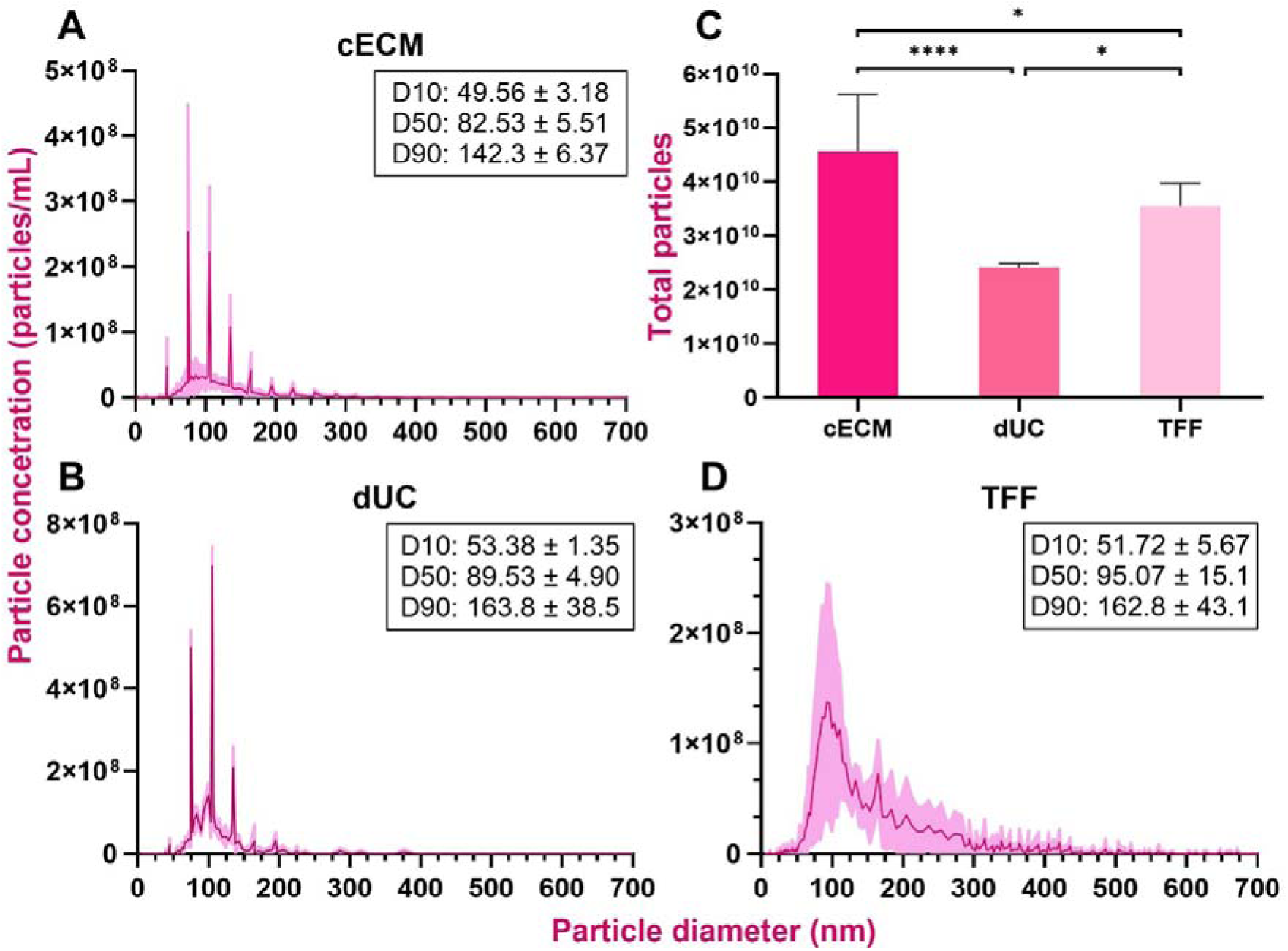
Particle size distribution of *Schistosoma mansoni* egg-derived nonvesicular extracellular particles (eggNPs) from (A) clarified egg conditioned media (cECM) and the extracellular nanoparticles that were isolated and separated using (B) differential ultracentrifugation (dUC) or (D) tangential flow filtration (TFF). Data illustrate the average distribution of particles based on 3 x 60 second videos per sample using nanoparticle tracking analysis. The mean concentration is the red line, while the standard deviation is shown in pink. The boxes containing the D10, D50, and D90 correspond to the percentage of particles that are below the estimated particle size. TFF. n = 6 replicates from 2 biological replicates for all samples. Groups were compared using a one-way ANOVA with Tukey’s multiple comparison test *p = 0.0418 (cECM vs TFF); *p = 0.0274 (dUC vs TFF); **** = 0.0001. (D) Total particle numbers recorded in cECM and eggNPs purified using dUC or TFF. Groups were compared using one-way ANOVA with Tukey’s multiple comparison test. **** <0.0010; ** < 0.005. Results shown as mean ± SD.

### TFF resulted in the lowest degree of contamination

Despite exhibiting similar final particle concentration to dUC eggNPs, TFF recovered significantly more particles (76.19 ± 7.62%; p = 0.0274) than dUC, which showed significant variability (38.90 ± 30.78%) (**Figure 6D-E)**. In addition, both methods resulted in significantly less total particles than cECM (dUC: p < 0.0001; TFF: p = 0.0274). To evaluate the purity of eggNPs from each separation protocol, the ratio of particles to protein concentration was calculated as previously described (Equation 1) ^56^. The protein concentration for eggNPs was quantified using Qubit^TM^, MicroBCA^TM^, and NanoDrop 2000 (Thermo Fisher Scientific). Significant variability was observed among the protein quantification protocols; however, the Qubit protein assay gave the most consistent measurements (**Supplementary** Figure 2). Therefore, all purity calculations were undertaken using Qubit assay values. The TFF enrichment resulted in the highest particle purity (1.35 x 10^8^ ± 2.88 x 10^7^ particles/µg protein), which was significantly higher than both cECM (6.88 x 10^5^ ± 3.68 x 10^5^ particles/ µg protein; p < 0.0001) and dUC (9.09 x 10^7^ ± 3.77 x 10^6^ particles/ µg protein; p = 0.0048) (**Figure 7A**). The ratio of protein (ng) per egg was also calculated as a measure of particle purity (Equation 2). Like above, TFF-isolated eggNPs (1.82 ± 0.18 ng protein/egg) exhibited a significantly higher purity compared to dUC eggNPs (0.57 ± 0.34 ng protein/egg; p < 0.0001). Significant variation was observed across protein quantification methods, limiting the confidence in protein-related purity calculations. We therefore calculated the ratio of particles per *S. mansoni* egg (**Figure 7C**; equation 3). As expected, TFF was indeed the superior eggNP purification method, with 1.79 x 10^5^ ± 1.91 x 10^4^ particles/egg compared the 5.52 x 10^4^ ± 3.2 x 10^4^ particles/egg in dUC eggNPs (p < 0.0001). In addition, all eggNP samples were analysed for endotoxin contamination. The highest concentration of endotoxin was found in eggNPs separated by dUC (5.57 EU/mL), while TFF gave rise to eggNPs with the lowest endotoxin contamination (0.22 EU/mL), further supporting TFF as the superior purification protocol.

**Figure 7:**
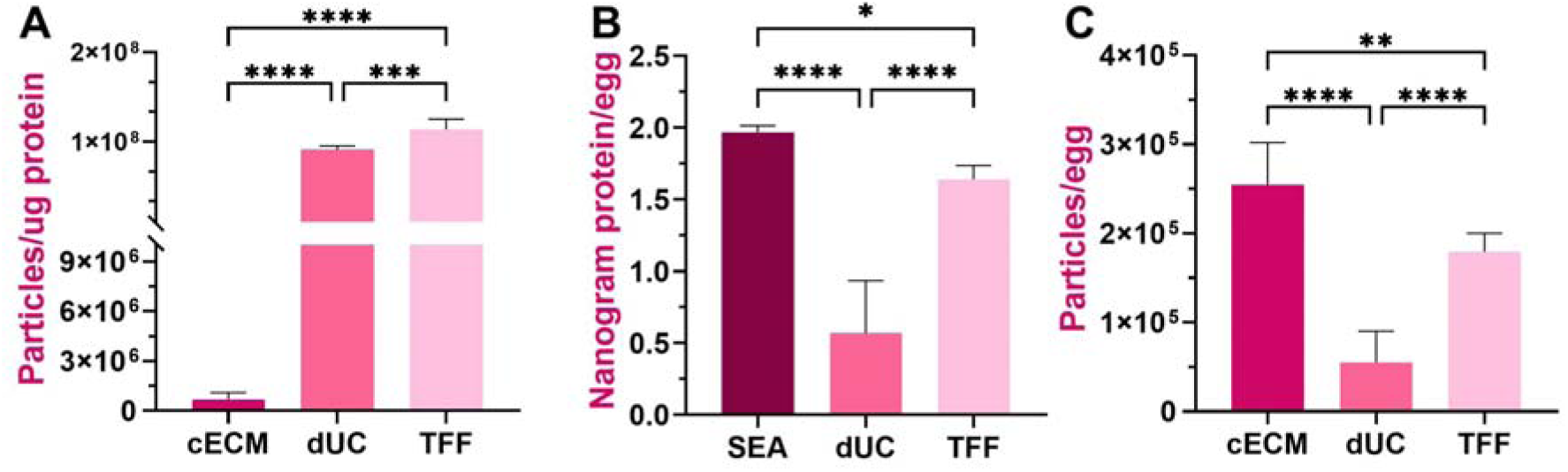
Characterising the purity of *Schistosoma mansoni* egg-derived nonvesicular extracellular particles (eggNPs). (A) Particle to protein ratio as an estimate of *S. mansoni* eggNPs purified using differential centrifugation (dUC) and tangential flow filtration (TFF) compared to clarified egg-conditioned media (cECM). The use of TFF gave the highest particle/protein ratio, 1.35 x 10^8^ ± 2.88 x 10^7^ particles/μg protein, indicating that these eggNPs have the lowest protein contamination. (B) Comparison of total protein (ng) per egg in *S. mansoni* soluble egg antigen (SEA), cECM, and eggNPs purified using dUC and TFF. TFF gave the highest protein (ng) per egg ratio of 1.82 ± 0.18 ng protein/egg (C) Comparison of total particles per *S. mansoni* egg in cECM, dUC eggNPs, and TFF eggNPs. TFF once again gave the highest purity particles with 1.79 x 10^5^ ± 1.91 x 10^4^ particles/egg. All groups were compared using one-way ANOVA with Tukey’s multiple comparison test. P values: ** < 0.005; **** < 0.0001. Results shown as mean ± SD.

### *S. mansoni* eggNPs contain enriched SEA-derived proteins

Silver stain was undertaken to visualise the protein profiles for eggNPs between separation methods and media controls (**Figure 8A)**. The protein profile for dUC and TFF-purified eggNPs were similar, with media-specific bands at 30kDa, 55kDa, and 60 kDa, and a number of distinct protein bands. However, in addition to the seven enriched bands at 10, 11, 13, 25, 40, 45, and 50 kDa seen in dUC eggNPs, TFF eggNPs exhibited an additional two low intensity bands at 14 and 17 kDa, which were not observed in the media control samples. Moreover, these bands were also observed in the protein profile of *S. mansoni* SEA, further supporting their egg-specific origin (**Figure 8B**). Western blotting (WB) with established EV-specific markers is an essential piece of evidence required by the MISEV2023. However, current available markers, such as heat-shock protein 90 (HSP90), are specific to mammalian cell-derived EVs. In addition, HSP90 was found to be present in both control media (RPMI-1640 GlutaMax) and cECM and were subsequently partially or fully removed during dUC and TFF, respectively (**Figure 7D**). Therefore, HSP90 could potentially be useful as purity marker for eggNPs. There is currently only one established marker for *S. mansoni* EVs, tetraspanin 2/5B (TSP-2), which is a rabbit polyclonal antibody raised against a conserved region of the hookworm aspartic protease that recognises *S. mansoni* TSP-2 (kindly provided by Professor Alex Loukas and Darren Pickering, James Cook University, Australia). TSP-2/5B was detected in both dUC and TFF eggNPs but not in cECM or control media, suggesting this protein was enriched alongside eggNPs during purification.

**Figure 8:**
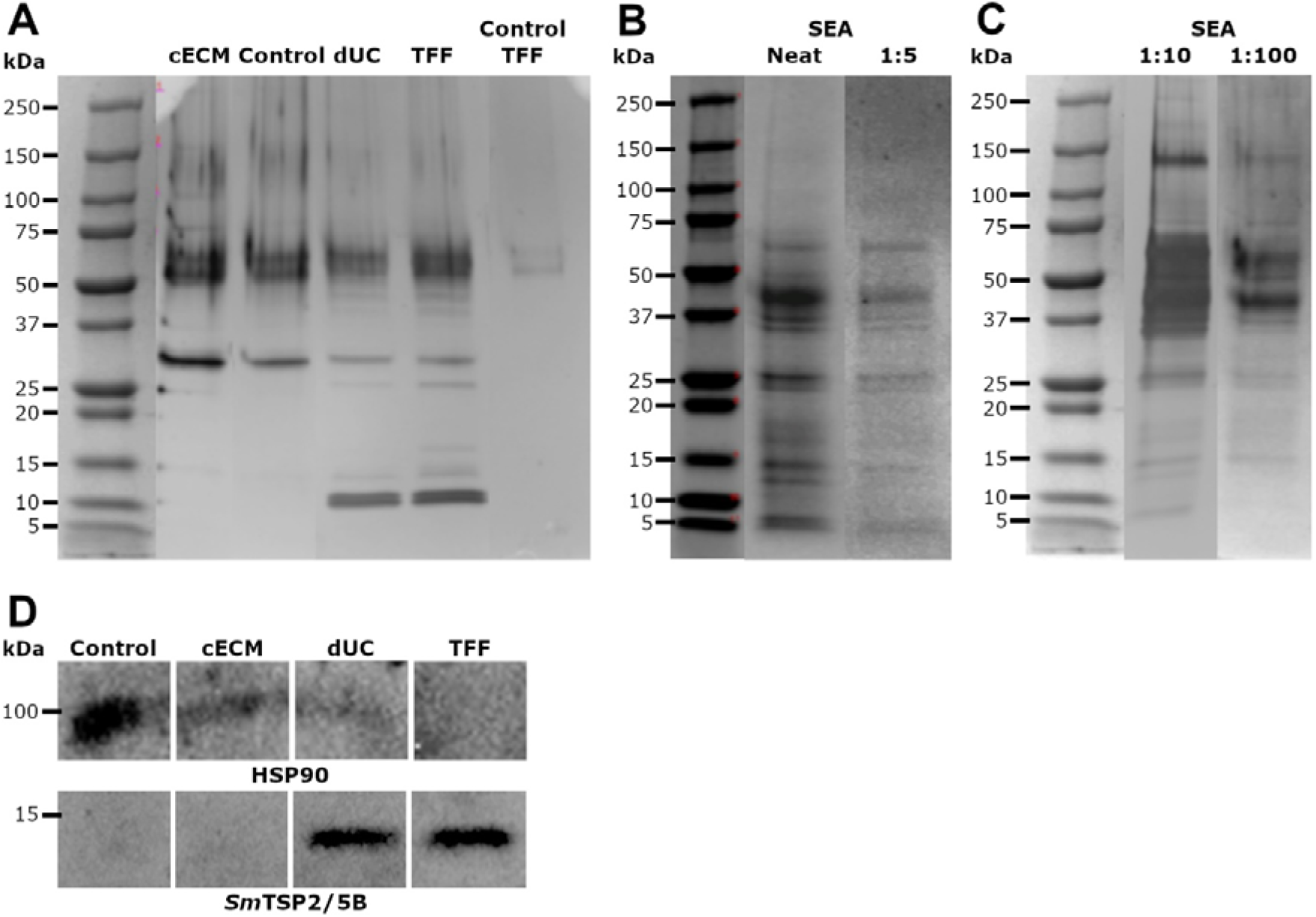
Characterising the protein profiles of *Schistosoma mansoni* egg-derived nonvesicular extracellular particles (eggNPs) (A) Silver stain profile of denatured clarified egg conditioned media (cECM), control media, differential ultracentrifugation (dUC) or tangential flow filtration (TFF) isolated eggNPs, and TFF media control. 500-1000 ng of protein was loaded onto the gel for each sample. (B) Coomassie blue stain profile of denatured soluble egg antigen (SEA) (Neat and 1:5). 2 and 10 µg of protein was loaded into the gel for the neat and 1:5 dilution respectively. (C) Silver stain profile of denatured diluted SEA (1:10 and 1:100). 0.1 and 1 ng of protein was loaded into the gel for the 1:10 and 1:100 dilution respectively. (D) Western blot detection of heat shock protein 90 (HSP90) and *S. mansoni* tetraspanin-2/5B in clarified egg-conditioned media. 1 - 2 µg of protein was loaded onto the gel for each sample.

## Discussion

This study initially aimed to develop a novel scalable method for the isolation of EVs from *S. mansoni* eggs and characterise their size, native structure, and proteomic content. Concerted efforts were made to purify EVs using dUC, and TFF. NTA, western blotting, and negative staining TEM suggested isolation was successful, and suggested the presence of vesicular particles sized between 50-150nm. However, cryoTEM imaging to characterise the native structure indicated that no vesicular nanoparticles were present in any preparation. Instead, we report the successful separation and identification of a novel population of 5 – 30 nm diameter NVEPs. These NVEPs were not present in control samples, indicating they are *S. mansoni* egg-derived (eggNPs).

CryoTEM imaging indicated that eggNPs separated by dUC were present in the form of tight aggregates where each individual eggNP was difficult to distinguish from the next (**Figure 3A-B)**. This was in contrast to the TFF enriched eggNPs, where single particles were easier to distinguish within aggregates (**Figure 3D-E)**. The aggregation observed in dUC eggNPs is likely due to the high-speed centrifugation (high shear force) required for separation. In contrast, the consistently low pressures used in the TFF (<0.4 Bar, low-moderate shear rate) prevent globularisation, resulting in more reproducible eggNP purification. EggNPs are irregularly shaped and exhibit significant adhesion, particularly when preserved in cryopreservation buffer. The CryoTEM imaging supported this, showing increased aggregation of cryobuffer-stored eggNPs that were obscured by large quantities of aggregated (most likely sucrose-derived) particles (**Figure 5D-F**). It is also possible that this was caused by the eggNPs possessing a low surface charge (zeta potential, ZP), making them more likely to aggregate, which was supported by the observation that eggNPs in PBS also exhibited significant aggregation (**Figure 5A-C**). Further analysis of eggNP ZP in cryobuffer is required to confirm this. The adhesive quality of eggNPs was likely also influenced by the co-isolation of other potentially adhesive compounds such as IPSE-α1 and omega-1 from the eggs or media.

Comparison by NTA of eggNPs separated by dUC or TFF identified that while the final particle concentration was similar, the most effective separation method was TFF, with a particle recovery of 76.19 ± 7.62%. In contrast, dUC particle recovery was highly variable (38.90 ± 30.78%). This variability is likely due to the low abundance of eggNPs secreted in culture, suggesting there is a minimum starting particle concentration required for efficient eggNP recovery using dUC. This is again likely necessitated by the significant particle loss occurring during high-speed differential ultracentrifugation, resulting in eggNPs absorbing onto the tube walls, and globularising into tightly packed flocs in the pellet. The protein profile for dUC and TFF eggNPs was similar, with a number of eggNP-specific bands that were also present in the SEA protein profile, suggesting similar contents and purity. However, the significantly higher particle to protein ratio was observed in TFF eggNPs compared to dUC eggNPs supports TFF as the superior isolation method. In addition, the scalable nature of TFF makes it ideal for isolating such low abundance nanoparticles. The protein profiles of TFF and dUC eggNP also exhibited media-specific bands. In addition, the microtubule-like and filamentous contaminants were observed by cryoTEM in TFF samples only (**Figure 4)**. The addition of size exclusion chromatography may assist in further eggNP purification and enrichment, however, further experimentation with a large enough starting particle number (>5 x 10^10^ particles) is required to confirm this, presenting a significant technological hurdle.

Although TFF eggNPs exhibited significantly higher particle to protein ratio than dUC, significant variability was observed in protein concentration across Nanodrop, microBCA, and Qubit assays, limiting the confidence of this purity measure (**Table S2**). Typical protein quantification methods rely on the assumption that the proteins of interest behave similarly to bovine serum albumin (BSA). However, many proteins, such as ovalbumin and IgG, are intrinsically disordered and do not fold into well-defined globular 3D structures due to their highly variable amino acid composition ^59,60^. These intrinsically disordered proteins (IDPs) are usually enriched in glutamine, glycine, serine, lysine, glutamine, and proline, which promote disorder, and depleted in valine, leucine, cysteine, phenylalanine, tyrosine and tryptophan, which promote order ^60^. IDPs result in unexpected Abs280 values due to macromolecular contaminants and will hardly interact with the dyes used in colorimetric assays ^59,60^. It is also possible that eggNPs may exhibit high carbohydrate contents. *S. mansoni* eggs produce of myriad glycoconjugates, the carbohydrate components of which play essential roles in host immune regulation and parasite survival, where they may act as antigenic motifs and ligands for host carbohydrate-binding proteins ^61^. This makes them particularly problematic for Bradford and BCA assays, which will often overestimate the protein concentration ^59,60^. Elemental analysis, which measures the amount of carbon and nitrogen present per gram of protein, is the most accurate way to quantify IDPs, however, the cost, time, and equipment requirements are a significant hurdle ^60,62^. Finally, Qubit is a highly accurate alternative for globular proteins, however, as the protein unfolds, the error of Qubit increases ^60^. This is likely why SDS-PAGE of eggNPs showed clear enriched egg-derived protein bands despite the low protein concentration. Despite this, multiple unsuccessful attempts were made to characterise the observed protein bands using LC-MS proteomics. Due to their complexity, the stability of both unbound and bound IDPs can depend heavily on both temperature, buffer, pH, and freeze/thawing cycles. It is therefore possible that the temperature used to denature (95°C for 5 mins) samples for SDS-PAGE prior to LC/MS on proteins bands was detrimental to their downstream characterisation. In support of this hypothesis, when the SDS-PAGE was repeated with altered denaturation conditions (75°C, 5 mins), additional bands were visible in both dUC and TFF eggNPs (**Figure S3**).

One way to overcome protein quantification issues is the use of a protein standard reference, such as the amount of soluble protein per *S. mansoni* egg. However, there is currently no such standard reference published. We therefore purified SEA from a controlled number of *S. mansoni* eggs and quantified protein using BCA, Nanodrop2000, and Qubit. Similar to eggNPs, significant variation in protein quantity was observed across methods, further suggesting the presence of IDPs (**Table S3**). Using Qubit protein quantities, the nanograms (ng) of protein egg was calculated and compared that of dUC and TFF eggNPs. This provided a more reliable measure of protein quantities and indicated that TFF eggNPs contained significantly more enriched proteins than dUC eggNPs (**Figure 7B)**. To improve the confidence of these results, however, elemental analysis would be required. In the meantime, we have developed an additional measure of particle purity using the ratio of particles per *S. mansoni* egg (**Figure 7C**). Using this measure, we confirmed that TFF gave rise to eggNPs with significantly higher purity than dUC.

In addition to the above measure, the MISEV2023 recommends to confirm successful purification by immunoblotting with antibodies for an EV-specific proteins as a marker of enrichment and purification ^37^. We therefore used the *S. mansoni* EV-specific antibody for TSP-2 as a marker of enrichment, and HSP90, which is present in the egg-free media as a marker of purification success (**Figure 8D**). Tetraspanins (TSPs) are plasma membrane-bound proteins that have the potential for use as novel diagnostic biomarkers and vaccine targets for schistosomiasis due to their accessibility to the host immune system and protective roles against schistosomes ^63,64^. TSP-2 is expressed in the tegument of cercariae, schistosomula, and adults (and their tegumental vesicles) but not eggs, making its presence and enrichment in *S. mansoni* eggNPs surprising as no membrane bound structures were observed ^64–67^. However, TSP-2 mRNA has been found to be transcribed by *S. japonicum* eggs ^66^. In addition, a recent single-cell RNA sequencing study of *S. mansoni* miracidia identified populations of *tsp2^+^*expressing Delta/Phi stem cells and tegument cells ^63^. It is unlikely that TSP-2 was co-isolated with eggNPs due to the molecule size cut-off used for TFF of 300 kDa which would only concentrate particles that are 30 nm or higher. *Sm*TSP-2 is only 25 kDa, which corresponds to ∼2.5 nm, making it too small to be concentrated by TFF unless it was associated with a larger structure, such as eggNPs. It may still be possible that a small number of small EVs (30 - 50 nm) were also present among eggNPs, which would explain the presence of *Sm*TSP-2, however, we examined a multiple slides from each biological replicate with both negative staining TEM and cryoTEM and did not observe membrane structures. It is also important to note that the molecular size cut-off used (300 kDa) for TFF, which was initially selected to isolate EVs of 50-150nm, likely also resulted in the loss of eggNPs during purification. Therefore, future isolations of eggNPs would benefit from TFF using a <50 kDa cut-off to ensure all eggNPs are recovered. This would likely result in the co-isolation of other protein contaminants, resulting in the requirement of additional purification steps such as size exclusion chromatography.

To date, there are no studies focussing on schistosome-derived NVEPs; however, analysis of the negative staining TEM images in the four available papers focussing on schistosome egg-derived EVs suggests these crude samples may also contain populations of eggNPs. Given our significant struggles with imaging eggNPs by negative staining, and the fact that the available evidence of EVs does not adhere to the MISEV2023 guidelines, it is also possible that these studies did not actually purify EVs, but some other miRNA and protein carrying EP species instead ^32–35,37^. This is further supported by comparing negative staining TEM images of eggNPs, which exhibited structural similarities to previously isolated so-called egg-derived EVs (**Figure 9**) ^32–35,37^.

**Figure 9:**
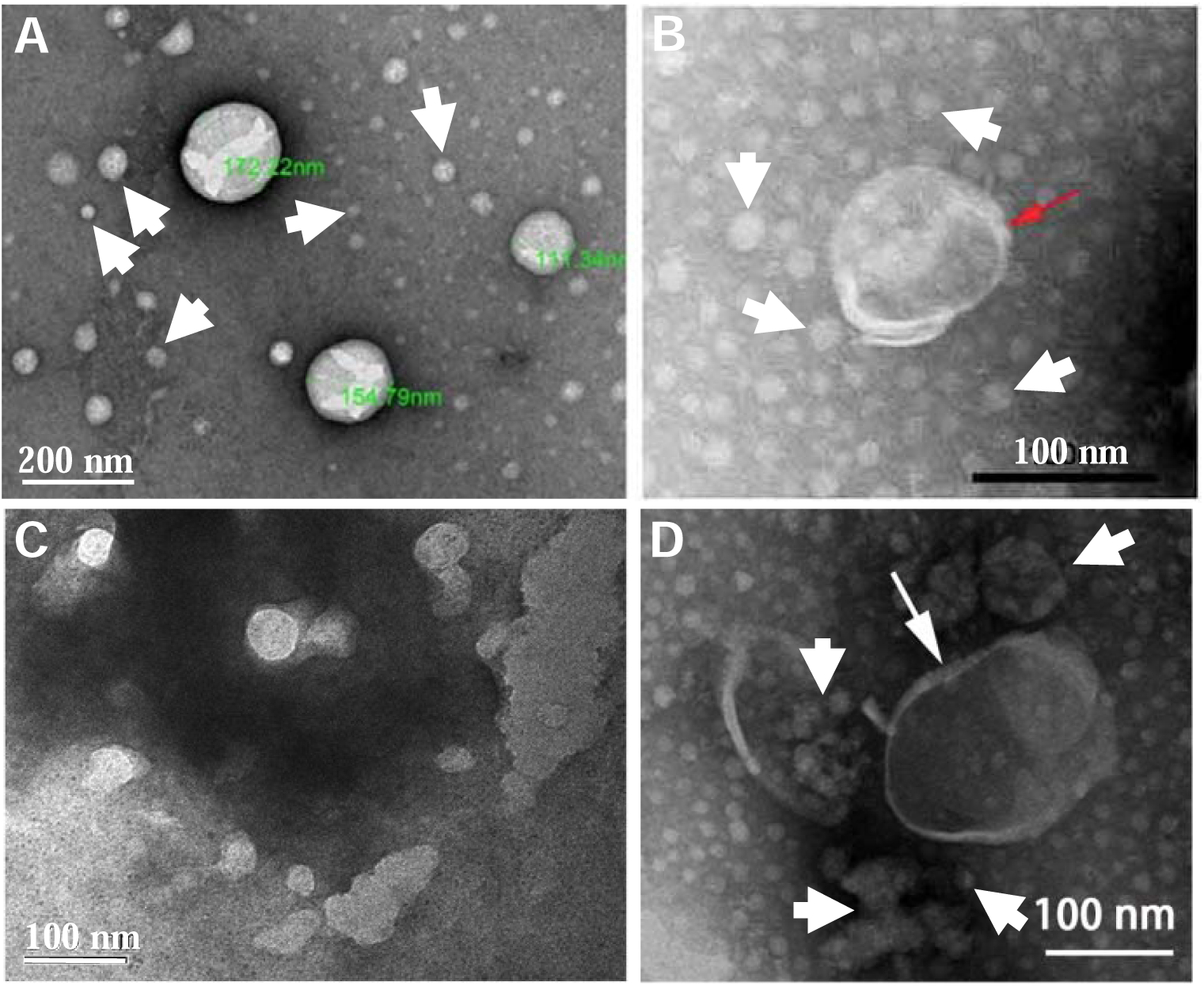
*Schistosoma* spp. egg-derived extracellular vesicles (EVs) – the current evidence is inconclusive. (A) Transmission electron microscopy (TEM) of “EVs” isolated from *S. mansoni* egg using differential ultracentrifugation dUC)^33^. (B, D) TEM of “EVs” isolated from *S. japonicum* eggs using dUC ^29,32^. (C) TEM characterisation of *S. japonicum* eggs using ExoQuick-TC Exosomes Precipitation Kit (System Bioscience) ^34^. The larger particles in images A-B and D exhibit limited resemblance to EVs, but are more likely to be densely packed aggregates formed during high-speed ultracentrifugation. Instead, it is possible that the smaller particles present behind these larger “EVs” (indicated by white arrows) correspond to nonvesicular extracellular particles like eggNPs. Cryogenic TEM is therefore required to confirm their structure and origin in native form. Image C is likely not of EVs at all but instead depicts the texture of the carbon-formvar coating on copper TEM grids.

The structure of *S. mansoni* eggs also likely impedes the release of such large and complex EVs. Schistosome eggs form through the polymerisation of tyrosine-rich eggshell precursor proteins (synthesised in the vitelline glands of the adult females), which are oxidised to *o*-quinones by tyrosinases ^68^. These *o*-quinones are subjected to nucleophilic attack by the lysine and histidine residues in the same or adjacent eggshell precursors, which leads to robust cross-link polymers ^68–70^. The largely impermeable eggshell is interrupted with complex nanoscale pores, the size of which is yet to be defined ^71,72^. These cribriform sieve-like pores consist of minute branching and anastomosing channels that are filled with fibrils and matrices similar to that of the outer envelope ^72^. Given their complex structure and the established resistance of schistosome eggs to penetration by fixatives, it is likely that these extra-miracidial layers obstruct the release of larger membrane bound-vesicles that may have been secreted by the tegument of the developing miracidia ^71,72^. The developing miracidia is surrounded by a dense acellular layer, known as the outer envelope, consisting of branching fibrillar structures that align closely with the shell and extend into the pores and a cellular inner layer ^71,72^. The Egg E/S products are thought to predominantly arise from the inner envelope, which must exit through the pores, significantly limiting the molecules that can be released and suggesting that any E/S molecules may require active secretion. Transmission electron microscope imaging of *S. japonicum* egg ultrastructure found an abundance of accumulated rosette bodies that resembled α-glycogen rosettes, (100-150nm) which were packaged into membrane-bound vacuoles known as Cheevers bodies ^71,72^. These appear to consist of smaller nanoparticles (10-30 nm) organised into rosettes, similar to dUC isolated eggNPs. However, the rosettes observed in whole eggs are significantly smaller than dUC eggNPs organised into rosette-like structures. In addition, this organised arrangement of eggNPs was not observed in TFF samples, further suggesting that the rosette structures formed due to high speed ultracentrifugation. The densely packed outer envelope may further impede EV biogenesis and secretion. Interestingly, imaging of *S. japonicum* eggs found granular material attached to the fibrils in the outer envelope have a similar size and structure to eggNPs ^71^. However, further imaging of egg ultrastructure at higher magnification is required to confirm this. Therefore, we hypothesise that *S. mansoni* eggs may have evolved specialised mechanisms for the secretion of EPs in the form of functional cargo-loaded NVEPs that contain miracidial tegument-derived proteins such as TSP-2.

## Acknowledgements

We would like to thank the following collaborators for their contributions to this research. A/Prof Laurence Macia and Dr Jian Tan from the University of Sydney Charles Perkins Centre kindly undertook all nanoparticle tracking analysis of EP samples isolated during this study. Dr Na’ama Koifman and Dr Matthias Floetenmeyer from the Microscopy Australia Facility at the Centre for Microscopy and Microanalysis (CMM) provided essential training in, and assistance with imaging eggNPs by cryogentic transmission electron microscopy. We further acknowledge Professor Alex Loukas and Darren Pickering, from James Cook University Centre for Biodiscovery and Molecular Development of Therapeutics for their kind donations of polyclonal sera against *S. mansoni* tetraspanin-2.

This research was undertaken under the guidance guidance of Vale Professor Don McManus, a distinguished and internationally recognised parasitologist, whose sudden passing in 2023 left his lab family and mentees with a significant loss. We would like to honour his memory by continuing his work and acknowledging his active contribution to the present manuscript.

SN is supported by a grant from the Children’s Hospital Foundation (RCP10317).

## Author Contributions

M.J.R., S.N., C.A.G., A.A. and J.B concieved and designed experiments. M.G.D. and M.R. performed all parasite material production and collection. M.J.R., with the guideance of J.B., undertook the majority of protocol optomisation and data acquisition and M.R. performed analyses from all experiments. N.C., A.A., and Y.L. contributed to the performance and anlyses of experiments. M.R. wrote the manuscript with the assistance and guidance of S.N., C.G., J.B., and M.K.J., who provided intellectual guidance and feedback.

