## Supplementary data for "Cracking the egg: Isolation and discovery of distinct nonvesicular extracellular particles from *Schistosoma mansoni* eggs"

### **Appendix 1 – Supplementary material**

**Figure S1:** Negative staining transmission electron micrographs of Schistosoma mansoni egg-derived nonvesicular extracellular particles (eggNPs) purified using differential ultracentrifugation (dUC) or tangential flow filtration (TFF). (A-B) dUC separated eggNPs imaged at 40,000x magnification. TFF separated eggNPs at 40,000x (C) and 60,000x magnification. Scale bars = 100 nm.

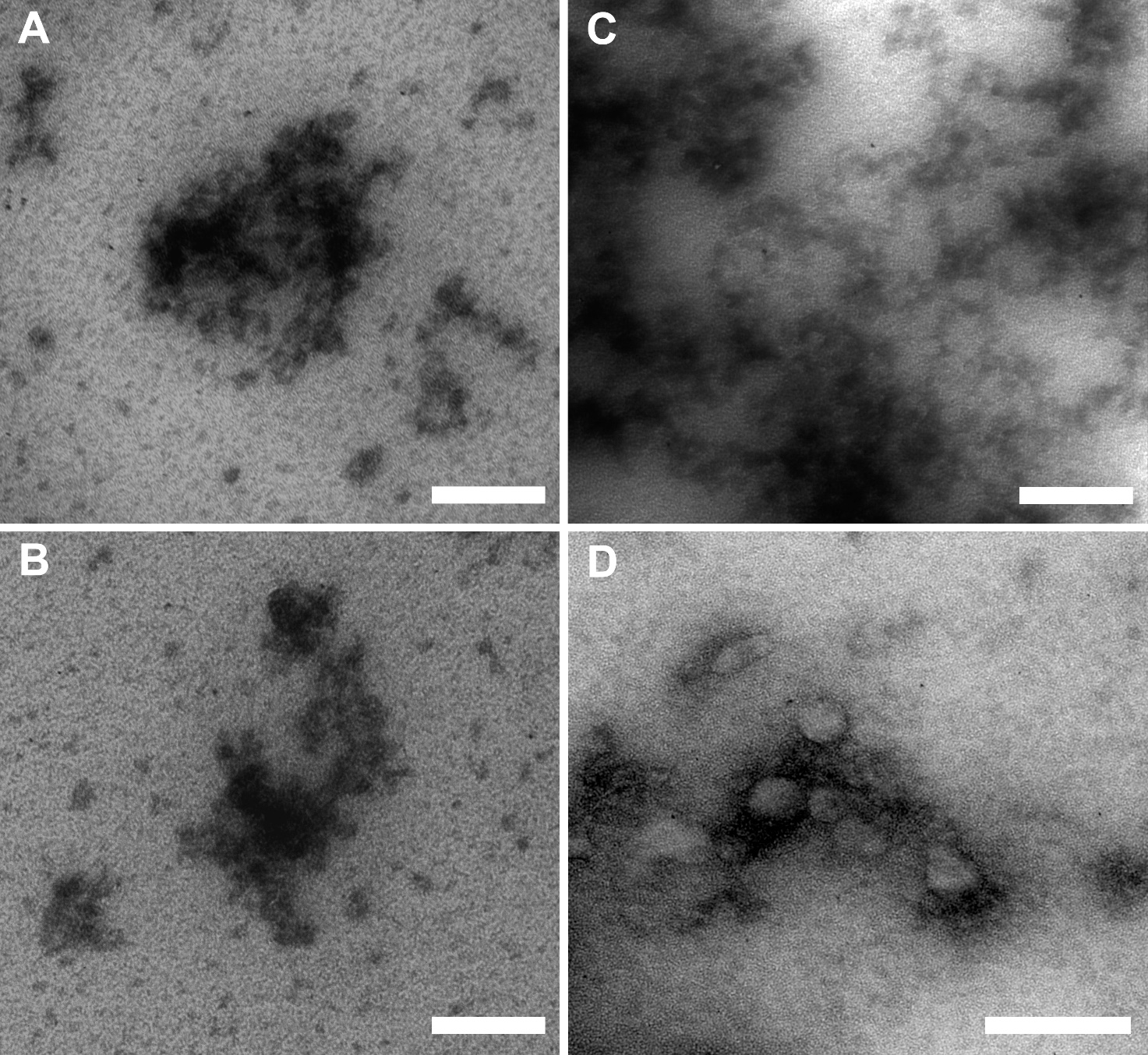

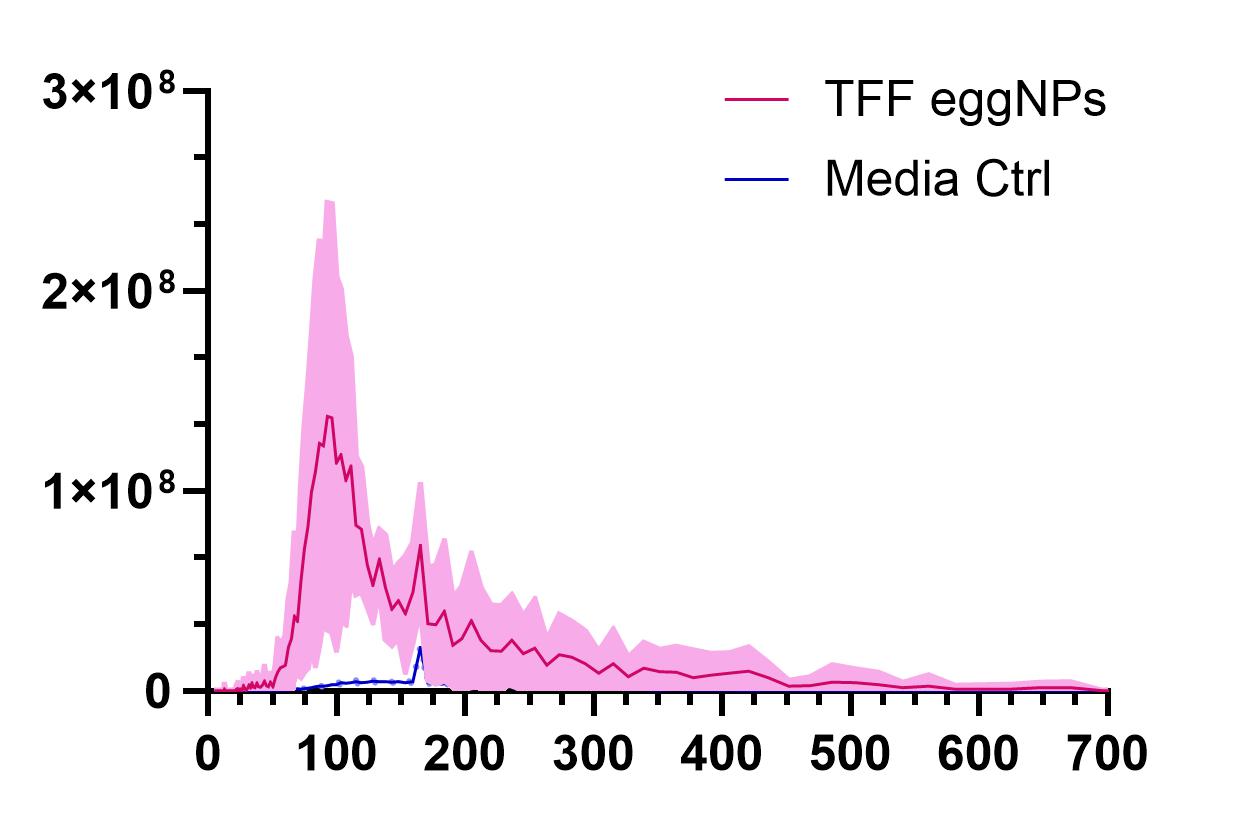

**Figure S2:** Particle size distribution of *Schistosoma mansoni* egg-derived nonvesicular extracellular particles (eggNPs) that were isolated and separated using tangential flow filtration (TFF) (pink line) compared to TFF processed control media (blue line). Data illustrate the average distribution of particles based on 3 x 60 second videos per sample using nanoparticle tracking analysis. The mean concentration is the red/blue line, while the standard deviation is shown in pink/light blue. n = 6 and n= 4 replicates for eggNP and media control samples respectively.

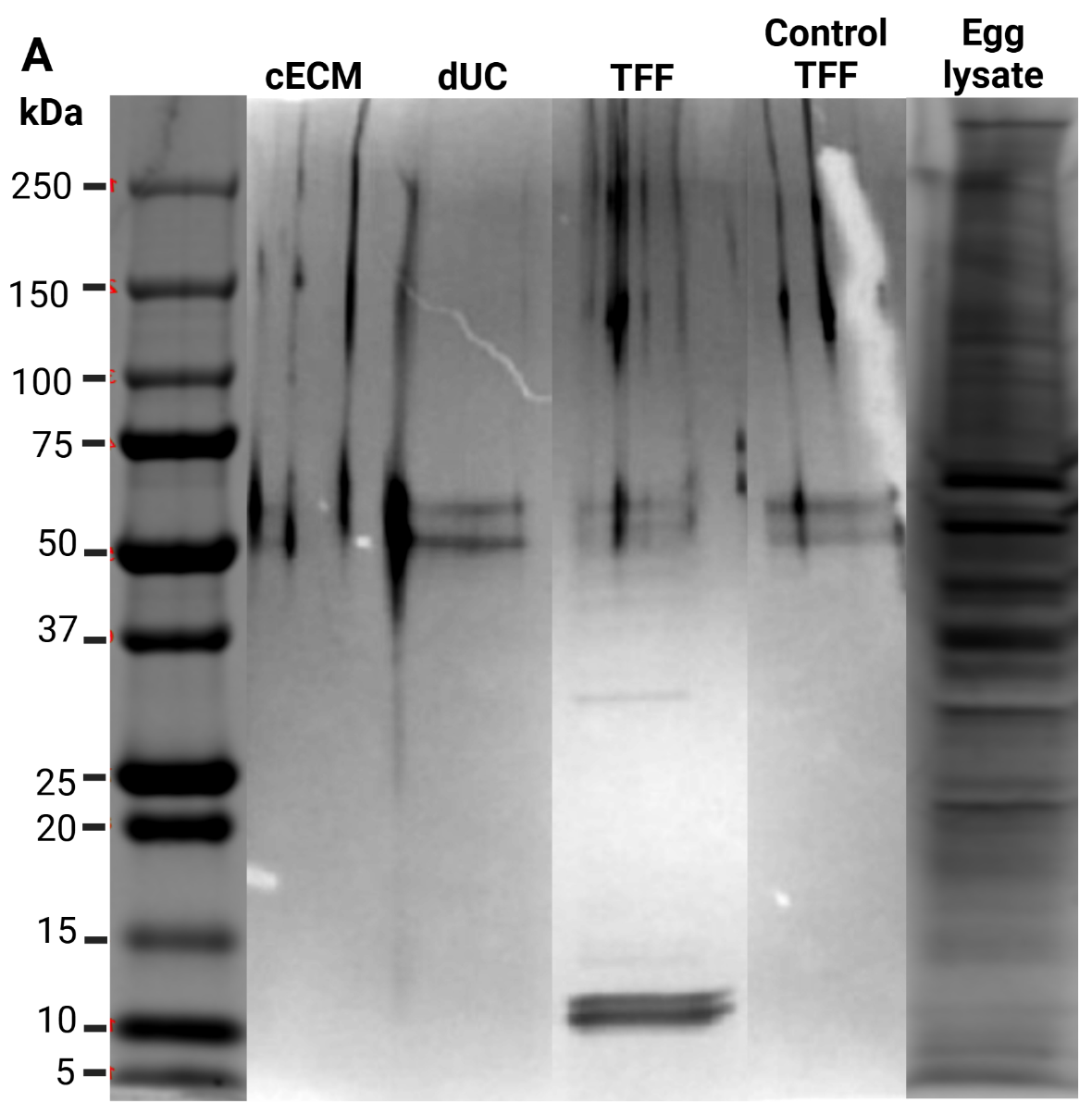

**Figure S3:** Silver stain profile of denatured Schistosoma mansoni clarified egg-conditioned media (cECM), differential centrifugation (dUC) or tangential flow filtration (TFF) isolated eggNPs, TFF media control end egg lysate. Samples were denatured at 95℃ for 5 minutes. 1100 ng of cECM protein was loaded onto the gel, while 722 ng, 990ng, 1533, and 1778ng, was loaded onto the gel for dUC, TFF, control TFF, and egg lysate samples respectively.

**Table S1*:*** Overview of details from experiments on the isolation, characterisation, and functional analysis of nonvesicular extracellular particles isolated from *Schistosoma mansoni* eggs (eggNPs) using differential centrifugation and tangential flow filtration as prescribed by MISEV2023 (Welsh *et al*., 2024) and the community-led roadmap by White and colleagues (2023).

| **Criteria for reporting** | | **Suggested standard** | **Reported** | | **Comments** |
| --- | --- | --- | --- | --- | --- |
| **Helminths** | | | | | |
| Helminth species including strain, genotype, drug specificity if applicable | | Mandatory | *Schistosoma mansoni* | | Strain: Navel Medical Researchg Institute (NMRI) |
| Life stage of helminth | | Mandatory | Eggs | | Isolated from liver of infected mice |
| Number, weight, sex and/or size or volume of worms | | Mandatory if applicable | 2,400,000 | | *In vitro* cultured eggs |
| Washing buffer or media used (for washes?) | | Mandatory | 1mM PBS (0.22 um filtered) | | Thermofisher (61870036) |
| **Host** | | | | | |
| Host species (and strain) used and origin when not a laboratory reared host (field isolates) | | Mandatory | Swiss mice | |  |
| Host age and sex | | Mandatory | 7-9 weeks, female | |  |
| Host tissue from where the helminth is isolated (Blood, faeces, etc) and tissue weight | | Mandatory | Liver; ~0.25g per liver | | ~25,000 eggs per liver; ~1000 eggs per g of liver tissues |
| Treatment if any | | **Encouraged** | 120 - 150 cercariae per mouse | | Cercariae shed from S. mansoni, strain Navel Medical Research Institute (NMRI), infected Biomphalaria glabrata snails (BEI Resources, NR-21962). |
| Number of host animals used | | **Encouraged** | 90 mice total | | dUC: 30 mice; TFF: 60 mice |
| **Helminth cultivation** | | | | | |
| Culture length | | Mandatory | 3 days | | Media collected and replenished every 24 hours |
| Composition of culture media | | Mandatory | PMI 1640 Medium, GlutaMAX™ Supplement; 1% Penicillan/Streptomyosin | | Media: ThermoFisher, 61870127; P/S: ThermoFisher, 15140122 |
| Volume of culture media | | Encouraged | dUC: 600 mL; TFF: 3000 mL | |  |
| Number of eggs per ml of media | | Mandatory if applicable | 1000-1500 eggs/mL | |  |
| Incubation temperature and % CO2 | | Mandatory | 37 degrees; 5% CO2 | |  |
| Parasite viability - description of viability assay performed (e.g. motility assay, visual inspection) | | Mandatory if applicable | Trypan blue viability staining and visual inspection by light microscopy | |  |
| Number of parasite pre-incubation washes and time | | Mandatory | Infected livers washed three times in sterile 1 mM PBS | | Eggs isolated from infected livers and by percoll gradient and washed in 1mM PBS before culture |
| Depletion of culture media of EVs if applicable (e.g. FBS used) | | Encouraged | N/A | |  |
| Biological contamination test (mycoplasma, LPS, ES agar plating, other) | | Encouraged | Limulus amoebocyte lysate (LAL) Assay | | Detection of viable and non-viable Gram-negative bacteria |
| **EP separation and QC** | | | | | |
| ES collection timepoints and storage | | Mandatory | Every 24 hours; placed into 25-30 mL aliquots, snap frozen in liquid N2, and stored -80 degrees | |  |
| EP storage details | | Mandatory | 30 - 500 uL aliquots, snap frozen in liquid N2, and stored -80 degrees | | Buffers: 1mM PBS and 1mM cryopreservation buffer (1 mM PBS, 5% sucrose, 50mM Tris, 2mM MgCl2) |
| EP separation technique used and details (details for reporting described in main text for each technique) | | Mandatory | Differential Ultracentrifugation OR Tangential Flow Filtration | |  |
| Details of low speed differential centrifugation and/or filtration performed to pre-clear ES of eggs and debris | | Mandatory | 1. 800g, 5 mins, 25 degrees and collect SN | |  |
|  |  |  | 2. Filter 0.22 um; | |  |
|  |  |  | 3. 3500g, 30 mins, 4 degrees and collect SN | |  |
| Upload of data in Vesiclepedia and other community-based open source repositories (PRIDE for proteomics data) | | Encouraged | N/A | |  |
| EVs number per worm, weight or size | | Encouraged | dUC | 1.79 x 10^5^ ± 1.91 x 10^4^ |  |
|  |  |  | TFF | 5.52 x 10^4^ ± 3.2 x 10^4^ |  |
| Reporting common residual host contaminants (i.e. host-albumin) | | Encouraged | GAPDH, HSP90 | | Media contaminants |
| Vesicle purity - particles/ug protein | Particles/µg protein | Encouraged | dUC | 6.88 x 10^5^ ± 3.68 x 10^5^ | Unit: total particles/ug protein |
|  |  |  | TFF | 1.35 x 10^8^ ± 2.88 x 10^7^ |  |
|  | Protein (ng)/egg | Optional | dUC | 1.82 ± 0.18 | Unit: total protein (ng)/egg |
|  |  |  | TFF | 0.57 ± 0.34 |  |
| **Functional studies of helminth EVs** | | | | | |
| Measurement of number of particles and /or size distribution and/or visualisation technique used (TEM, cryoEM) | | Mandatory if applicable | Nanoparticle tracking analysis (NTA), negative staining transmission electron microscopy (TEM); cryogenic TEM; | |  |
| Protein/lipid/glycan quantification | | Mandatory if applicable | microBCA, Qubit, Nanodrop2000 | |  |
| Proteomic analysis of specific EP or EP-enriched markers if available | | Encouraged | Western blot of eggNPs using polyclonal anti-*S. mansoni*-TSP2/5B antibody | | Raised in rabbits |
| Nucleic acid assessment and description of method used | | Encouraged | Small RNA sequencing | | TRIzol total RNA purification |
| EP labelling method | | Mandatory | N/A | |  |
| Controls of EV labelling and how they were obtained | | Mandatory | N/A | |  |
| Functional studies of eggNPs | | Mandatory | Type 1 and type 2 conventional dendritic cells cultured with eggNPs (14,000 eggNPs/cell) or in a transwell system with live eggs (500 egg/cell) | | Only undertaken on TFF eggNPs; Small RNA sequencing and bulk RNA sequencing |
| Controls used for functional studies and how they were obtained | | Mandatory | Type 1 and type 2 conventional dendritic cells cultured without eggNPs or in a transwell system without live eggs | |  |
| Normalisation of EVs for functional studies (i.e. number of vesicles/ml, number vesicles/worm) | | Mandatory | Total number of particles; Particles/mL; Particle recovery (%); Particles/ug protein; Protein (ng)/egg Particles/egg | |  |

**Table S2:** Comparison of Qubit, microBCA, and Nanodrop2000 for the quantification of protein in soluble egg antigen (SEA) isolated for *Schistosoma mansoni* eggs.

| Method | MicroBCA (µg/ml) | | Nanodrop (µg/ml) | | Qubit (µg/ml) | |
| --- | --- | --- | --- | --- | --- | --- |
| Sample | **Average** | **SD** | **Average** | **SD** | **Average** | **SD** |
| SEA replicate 1 | 508.7 | 6.47 | 3340 | 7.12 | 876.5 | 1.00 |
| SEA 2 replicate 2 | 604.6 | 6.40 | 3695 | 33.5 | 927.5 | 6.19 |
| SEA 3 replicate 3 | 637.18 | 3.94 | 3695 | 33.5 | 892.5 | 3.42 |

**Table S3:** Comparison of Qubit, microBCA, and Nanodrop2000 for the quantification of protein in nonvesicular extracellular particles, which were isolated for Schistosoma mansoni eggs (eggNPs) using differential ultracentrifugation (dUC) tangential flow filtration (TFF).

| Method | | MicroBCA (µg/ml) | | Nanodrop (µg/ml) | | Qubit (µg/ml) | |
| --- | --- | --- | --- | --- | --- | --- | --- |
| Sample | | **Average** | **SD** | **Average** | **SD** | **Average** | **SD** |
| BR1 eggNPs | **cECM** | 1328.13 | 39.64 | 490.67 | 6.55 | 116.67 | 0.47 |
|  | **dUC** | 4.70 | 0.00 | NM | NM | 40.03 | 8.76 |
|  | **TFF** | 13.47 | 17.27 | 20.90 | 7.18 | 22.00 | 1.63 |
| BR2 eggNPs | **cECM** | 1413.24 | 54.38 | 590.00 | 6.68 | 103.67 | 2.36 |
|  | **dUC** | 22.50 | 9.88 | 22.00 | 4.97 | 16.07 | 0.05 |
|  | **TFF** | 0.00 | 0.00 | 3.30 | 4.05 | 26.8 | 0.08 |
| Media ctrl | | 1545.32 | 10.45 | 449.67 | 18.21 | 126.50 | 3.50 |
| TFF media ctrl | | 3.35 | 0.61 | 9.33 | 3.09 | 35.87 | 1.80 |
| NM = Not measured | |  |  |  |  |  |  |
